# Deletion of serine palmitoyl transferase 2 in hepatocytes impairs ceramide/sphingomyelin balance, prevents obesity and leads to liver damage in mice

**DOI:** 10.1101/2021.09.16.460588

**Authors:** Justine Lallement, Ilyès Raho, Grégory Merlen, Dominique Rainteau, Mikael Croyal, Melody Schiffano, Nadim Kassis, Isabelle Doignon, Maud Soty, Floriane Lachkar, Michel Krempf, Fabienne Foufelle, Chloé Amouyal, Hervé Le Stunff, Christophe Magnan, Thierry Tordjmann, Céline Cruciani-Guglielmacci

## Abstract

Ceramides (Cer) have been shown as lipotoxic inducers, which disturb numerous cell signalling pathways especially insulin signalling pathway leading to metabolic disorders such as type 2 diabetes. In this study, we aimed to determine the role of *de novo* hepatic Cer synthesis on energy and liver homeostasis in mice. We generated mice lacking serine palmitoyltransferase 2 (*Sptlc2*), the rate limiting enzyme of Cer *de novo* synthesis, in hepatocytes.

Despite lower expression of hepatic *Sptlc2*, we observed an increased concentration of hepatic Cer, especially C16:0-Cer and C18:0-Cer associated with an increased neutral sphingomyelinase 2 expression, and a decreased sphingomyelin content in the liver. *Sptlc2*^ΔHep^ mice were protected against obesity induced by high fat diet. Bile acid (BA) hydrophobicity was drastically decreased in KO mice, and was associated with a defect in lipid absorption. In addition, an important increase of tauro-muricholic acid in BA pool composition was associated with a downregulation of the nuclear BA receptor FXR target genes. *Sptlc2* deficiency also enhanced glucose tolerance and attenuated hepatic glucose production. Finally, *Sptlc2* disruption promoted apoptosis, inflammation and progressive development of hepatic fibrosis worsening with age.

Our data suggest a compensatory mechanism to regulate hepatic Cer content from sphingomyelin hydrolysis, with deleterious impact on liver homeostasis. In addition, our results show the implication of hepatic sphingolipid modulation on BA metabolism and hepatic glucose production in an insulinin-dependent manner, which demonstrates the role of Cer in many metabolic functions still under-researched.

## 1. Introduction

Type 2 diabetes (T2D) affects both the quality of life and life expectancy of about 400 million people worldwide, and it is considered as a non-infectious pandemic due to the constant increase in the number of patients. The established disease is characterised by liver, fat and muscle insulin resistance, and by inadequate pancreatic beta-cells insulin secretion to counteract the insulin resistance.

A specific class of lipids, namely sphingolipids, and in particular ceramides (Cer), are proposed to be important mediators of both free-fatty acid (FFA)-induced β cell dysfunction and apoptosis, and FFA-induced insulin resistance in insulin target tissues (Poitout et Robertson 2008). Moreover, we previously showed that Cer and dihydroceramides could be relevant plasma biomarkers of T2D susceptibility (Wigger et al. 2017). At the cellular level, increased concentrations of Cer promote insulin resistance by inhibiting insulin signalling pathway, and Cer can regulate autophagy, reactive oxygen species production, cell proliferation, inflammation and apoptosis (Chaurasia et Summers 2015).

In mammals, three main pathways have been described to produce Cer. First, the *de novo* synthesis pathway starts on the cytoplasmic face of the endoplasmic reticulum (ER) with the condensation of a coenzyme A-linked fatty acid (typically palmitoyl-CoA) and L-Serine by serine palmitoyltransferase (SPT) to form 3-ketosphinganine. Three different subunits of SPT called SPTLC1, SPTLC2, and SPTLC3, more recently identified (Hornemann et al. 2006) have been described. SPTLC1 could act as a dimer with SPTLC2 and/or SPTLC3, which carry out the PLP (pyridoxal 5-phosphate) binding motif, essential for the catalytic activity (Hornemann, Wei, et von Eckardstein 2007). Then, Cer are transported to the Golgi apparatus to be metabolized into more complex sphingolipids such as glucosylceramides and sphingomyelin (SM). Second, the catabolic sphingomyelinase pathway leads to the degradation of SM into Cer by sphingomyelinases and takes place at different sub-cellular localizations (plasma membrane, mitochondria, lysosome and Golgi) (Insausti-Urkia et al. 2020). The third pathway is called the “salvage pathway” from late endosomes/lysosomes, which generate Cer from the breakdown of complex sphingolipids (Kitatani, Idkowiak-Baldys, et Hannun 2008). Both, in salvage pathway and *de novo* synthesis pathway, Cer are produced by ceramide synthase (CerS) through N-acylation of a sphingoid base. There are six different isoforms of CerS which exhibit specificities for the chain length of the added acyl-CoA leading to the production of different Cer species (Wattenberg 2018). Among the different Cer species, C18:0-Cer and C16:0-Cer have been pointed out as apoptosis and inflammatory inducers and play a key role in the development of insulin resistance and steatohepatitis development (Pewzner-Jung, Brenner, et al. 2010; Raichur et al. 2014).

The liver is a key metabolic organ, which governs body energy metabolism. In particular, the intrahepatic accumulation of lipids and, especially Cer, observed in NASH syndrome (nonalcoholic steatohepatitis) strongly correlates with the risk of T2D (Birkenfeld et Shulman 2014). In addition, it has been demonstrated that *in vivo* or *in vitro* inhibition of SPT, using myriocin, decreases Cer accumulation in plasma and liver and improves hepatic fibrosis, steatosis and insulin sensitivity (Holland et al. 2007; Kasumov et al. 2015; M. Jiang et al. 2019).

Hepatocytes, the major parenchymal cells in the liver, carry out many metabolic functions, including the production of bile acids (BA). BA are amphipathic molecules, which allow dietary lipid absorption and also act as signaling molecules through their action on nuclear receptors such as the Farnesoid X receptor (FXR) or G protein-coupled BA receptors, such as Takeda G protein-coupled receptor 5 (TGR5) or S1P receptor 2 (S1PR2) (Makishima et al. 1999; Studer et al. 2012). FXR is activated by chenodeoxycholic acid (CDCA) but inhibited by tauro-beta-muricholic acid (T-β-MCA), and regulates BA synthesis (Goodwin et al. 2000; Sayin et al. 2013). Numerous studies have demonstrated the role of FXR signaling in glucose homeostasis and especially in hepatic glucose production (Sun, Cai, et Gonzalez 2021). Thus, beyond their detergent properties, BA modification could trigger metabolic disorders in an endocrine way. Interestingly, inhibition of intestinal FXR signaling prevents obesity induced by high fat diet and metabolic disease such as insulin resistance and fatty liver (C. Jiang, Xie, Li, et al. 2015). Recently, *Smpd3* (sphingomyelin phosphodiesterase 3), a gene encoded for the neutral sphingomyelinase 2 (nSMase2) and *Sptlc2* gene, both involved in Cer production, have been identified as FXR target genes (Xie et al. 2017; Q. Wu et al. 2021). These results suggest the existence of a Cer/FXR/BA signaling axis, which could regulate glucose and lipid metabolism.

Because of a key role of hepatic sphingolipids production in aetiology of metabolic diseases we investigated the role of *de novo* Cer synthesis in the liver of mice fed with regular chow or HFD on energy homeostasis. Using the cre-lox system, we developed a mouse model to decrease Cer *de novo* synthesis in liver by targeting the rate-limiting enzyme, *Sptlc2*, in hepatocytes. Surprisingly, we found that a 60% decrease of liver *Sptlc2* expression led to the increase of liver Cer content at the expense of SM. In addition, hepatic SPTLC2 deletion impaired BA synthesis and export, associated with a defect in lipid absorption. Our results also showed that *Sptlc2*-deficient mice were protected against HFD-induced obesity and displayed impaired hepatic glucose production.

Altogether, we demonstrated that deletion of *Sptlc2* in hepatocytes induced Cer synthesis remodeling in favor of accumulation of deleterious C16:0-Cer and C18-Cer and was associated with strong alteration in BA homeostasis, decreased hepatic glucose production, and progressive liver fibrosis.

## 2. Results

### 2.1 Genetic deletion of *Sptlc2* in hepatocytes increases a selective ceramide content in the liver but decreases sphingomyelin content associated with an up-regulation of *nSmase2*

*Sptlc2*^lox/lox^ mice were crossed with heterozygous mice, which expressed the protein CRE driven by the albumin promoter (Figure 1 - figure supplement 1.A). *Sptlc2* hepatic mRNA level was reduced to 60% of WT level both in *Sptlc2*^ΔHep^ mice fed with regular chow (RC) or high fat diet (HFD) (Figure 1 - figure supplement 1.B), consistent with the fact that hepatocytes represent about 70% of all liver cells (Si-Tayeb, Lemaigre, et Duncan 2010). Surprisingly, in our genetic model, *Sptlc2* deficiency in hepatocytes does not induce ceramide (Cer) reduction in liver of RC mice (Figure 1.A) and even elicits hepatic Cer accumulation in mice fed HFD (Figure 1.B). Moreover, *Sptlc2* deletion modifies hepatic Cer species distribution. Thus, we showed C16:0-Cer, C18:0-Cer and C24:1-Cer accumulation and C22:0-Cer reduction in the liver of mice fed with RC or HFD (Figure 1.C and D). Quantification of Cer in isolated hepatocytes from *Sptlc2* ^ΔHep^ mice and their littermate controls confirmed the increase of hepatic C16:0-Cer and the decrease of C22:0-Cer (Figure 1 - figure supplement 1.C). Interestingly, *CerS5* and *CerS6*, which catalyze C16:0-Cer and C18:0-Cer synthesis are up regulated in liver of *Sptlc2* knock – down (KD) mice upon RC and HFD (Figure 1.E and 1.F).

**Figure 1.**
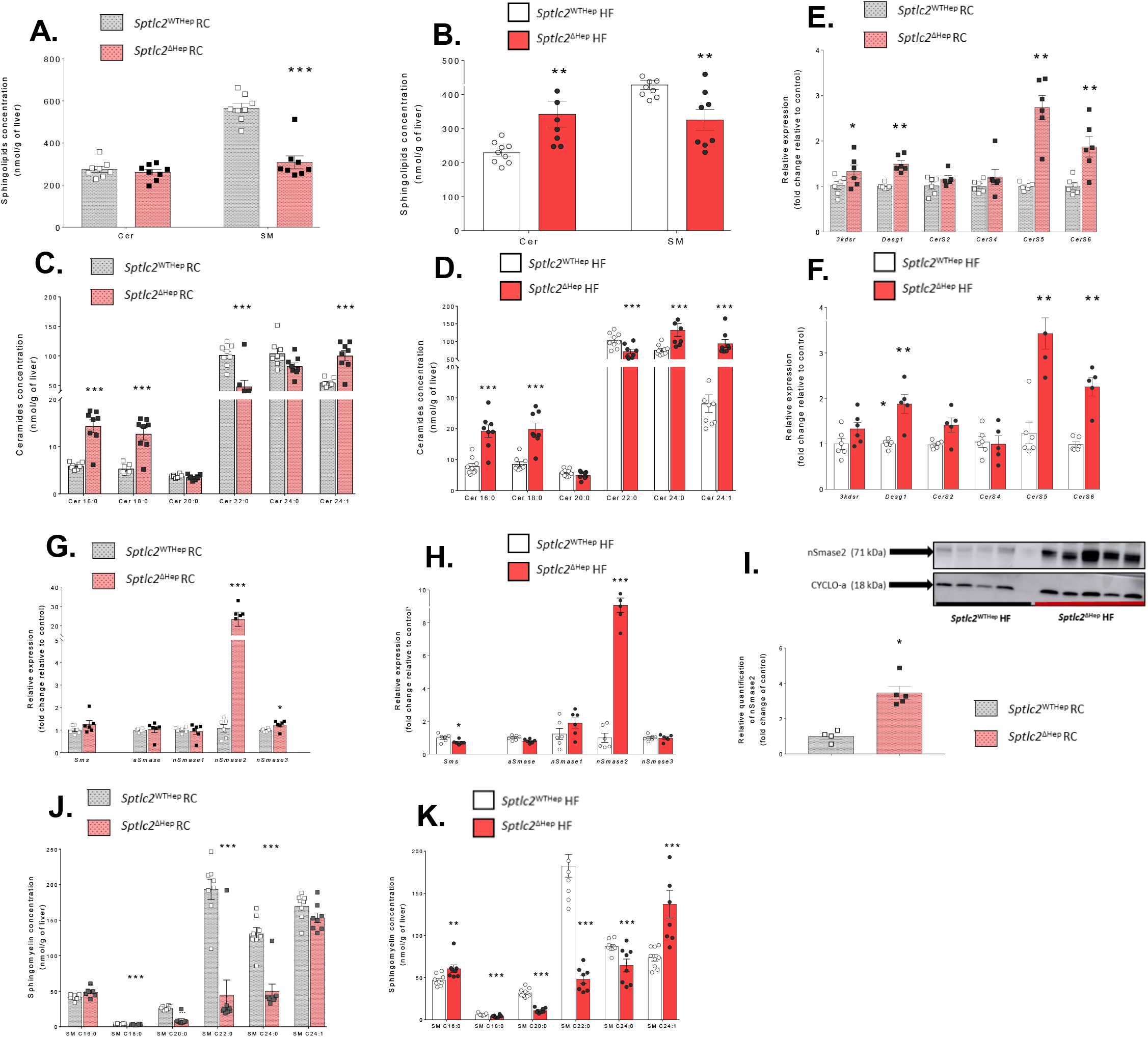
*Sptlc2* deficiency in hepatocytes modulates SM/Cer balance at the expense of Cer and induces *nSmase2* overexpression. (**A-B**) Quantification of SM and Cer content in the liver of regular chow (RC) mice and high fat diet (HFD) mice. (**C-D**) Ceramides species distribution according to acyl-chain length in the liver of RC and HFD mice. (**E-F**) Relative expression of genes involved in *de novo* ceramides synthesis in liver of *Sptlc2*^ΔHep^ mice relative to their littermate controls upon RC or HFD. (**G-H**) Relative expression of genes involved in sphingomyelinase pathway in liver of *Sptlc2*^ΔHep^ mice relative to their littermate controls upon RC or HFD. (**I**) Western blot analyses and quantification of nSMase2 and Cyclo-a in liver of RC mice. (**J-K**) SM species distribution according to acyl-chain length in the liver of RC and HFD mice. Analysis were performed in 4 month-old mice and HFD mice were fed with HFD for 2 months starting at 2 month-old; error bars represents SEM; n = 4 −10 mice per groups; *p < 0.05, **p < 0.01, ***p < 0.001 via Mann-Whitney U test. **RC**: Regular Chow; **HF**: High Fat; **Cer**: ceramide; **SM**: Sphingomyelin; **Sptlc1**: Serine Palmitoyltransferase long chain 1; **Sptlc2**: Serine Palmitoyltransferase long chain 2; **3kdsr**: 3-Dehydrosphinganine Reductase; **Degs1**: Delta-4-Desaturase Sphingolipid 1; **CerS2**: Ceramide Synthase 2; **CerS4**: Ceramide Synthase 4; **CerS5**: Ceramide Synthase 5; **CerS6**: Ceramide Synthase 6; **Sms**: Sphingomyelin Synthase; **aSMase**: acide Sphingomyelinase; **nSMase1**: neutral Sphingomyelinase 1; **nSMase2**: neutral Sphingomyelinase 2; **nSMase3**: neutral Sphingomyelinase 3; **Cyclo-a**: Cyclophilin-a.

Our data suggest that the sphingomyelinase (SMase) pathway from hydrolysis of sphingomyelin (SM) is over-activated to produce Cer. Indeed, sphingomyelin (SM) content was reduced in the liver of *Sptlc2*^ΔHep^ mice, at the expense of Cer (Figure 1.A and 1.B). SMase pathway takes place at different sub-cellular localizations and involves different SMases depending on the pH of the organelles (Insausti-Urkia et al. 2020). We measured the expression levels of *Sms* and *Smases* and we found that mRNA level of nSMase2 is dramatically elevated in the liver of KD mice, independently of the diet, compared to the littermate controls (Figure 1.G and 1.H). Moreover, up-regulation of nSMase2 protein levels was also detected by western blotting in the liver of *Sptlc2*^ΔHep^ mice (Figure 1.I). The analysis of SM species distribution reveals that the modification of SM profile in the liver is similar to this observed for Cer species (i.e increase of C16:0-SM and decrease of C22:0-SM) (Figure 1.J and 1.K) suggesting a remodeling of Cer species profile due to *CerS5* and *CerS6* up-regulation.

**Figure 1 - figure supplement 1.**
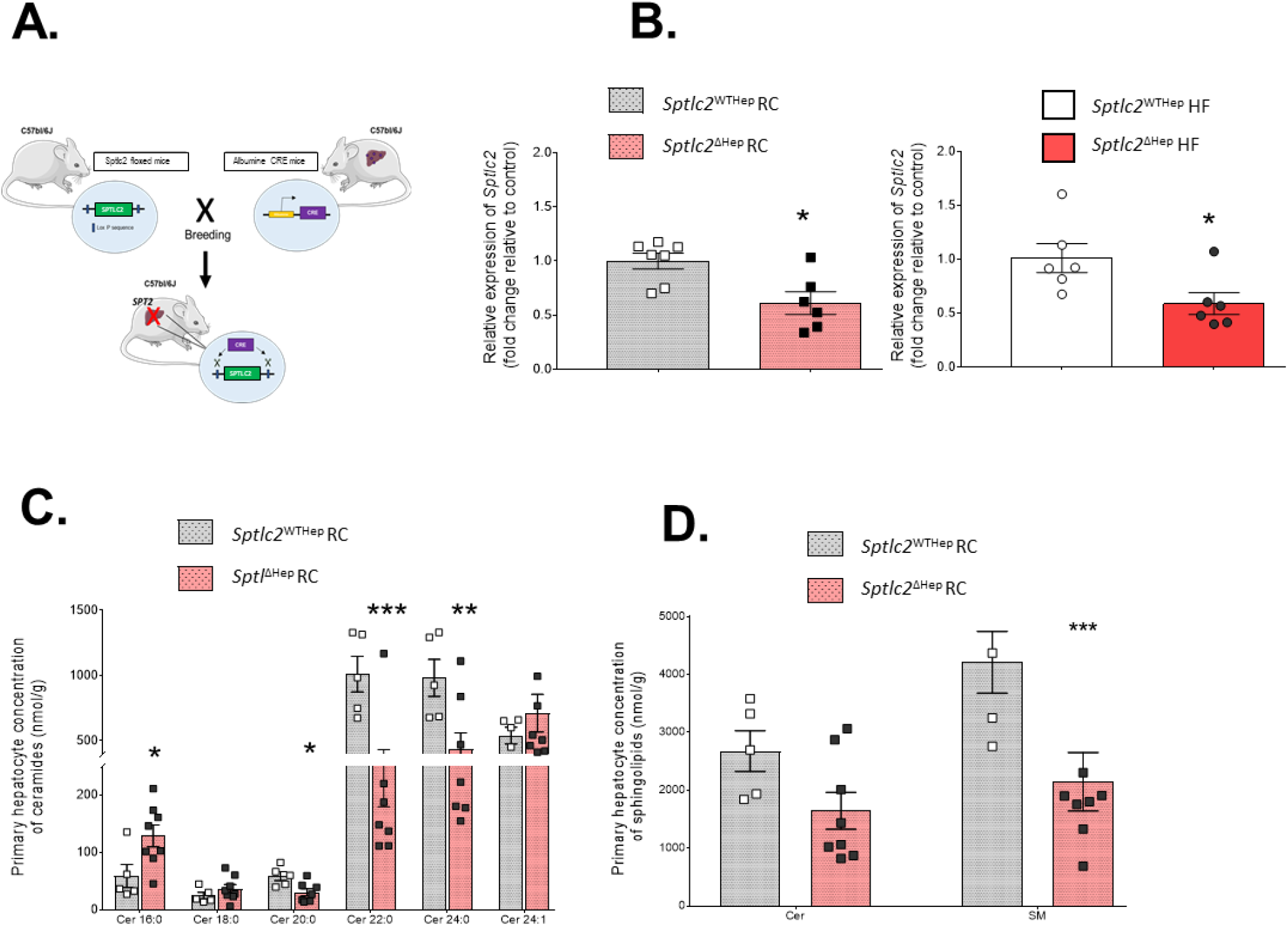
*Sptlc2*^ΔHep^ mice model and validation. **(A)** Schematic representation of breeding leading to *Sptlc2*^ΔHep^ mice. (**B**) Relative expression of *Sptlc2* in liver of *Sptlc2*^ΔHep^ mice and their control upon RC or HFD, data are expressed in fold change relative to control (**C**) Ceramides species distribution according to acyl-chain length in primary hepatocytes from *Sptlc2*^WTHep^ and *Sptlc2*^ΔHep^ mice upon RC (45 day-old) (**D**) Quantification of SM and Cer content in primary hepatocytes from *Sptlc2*^WTHep^ and *Sptlc2*^ΔHep^ mice upon RC (45 day-old). n = 4 - 7 mice per group; error bars represents SEM; *p < 0.05, **p < 0.01, ***p < 0.001 via Mann-Whitney U test or Student’s *t* when appropriate. **RC**: Regular Chow; **HF**: High Fat; **Cer**: ceramide; **SM**: Sphingomyelin; **Sptlc2**: Serine Palmitoyltransferase long chain 2

Cer and SM levels were also examined in the plasma and we found an increase of Cer and SM concentration in the plasma of mice lacking SPTLC2. Then, hepatic Cer and especially hepatic SM, substrate of nSMase 2, could be provided by plasma sphingolipids (Figure 1 - figure supplement 2).

Overall, these data suggest a compensatory mechanism through the SMase pathway to maintain an elevated Cer content in the liver despite the inhibition of *de novo* synthesis pathway.

**Figure 1 - figure supplement 2.**
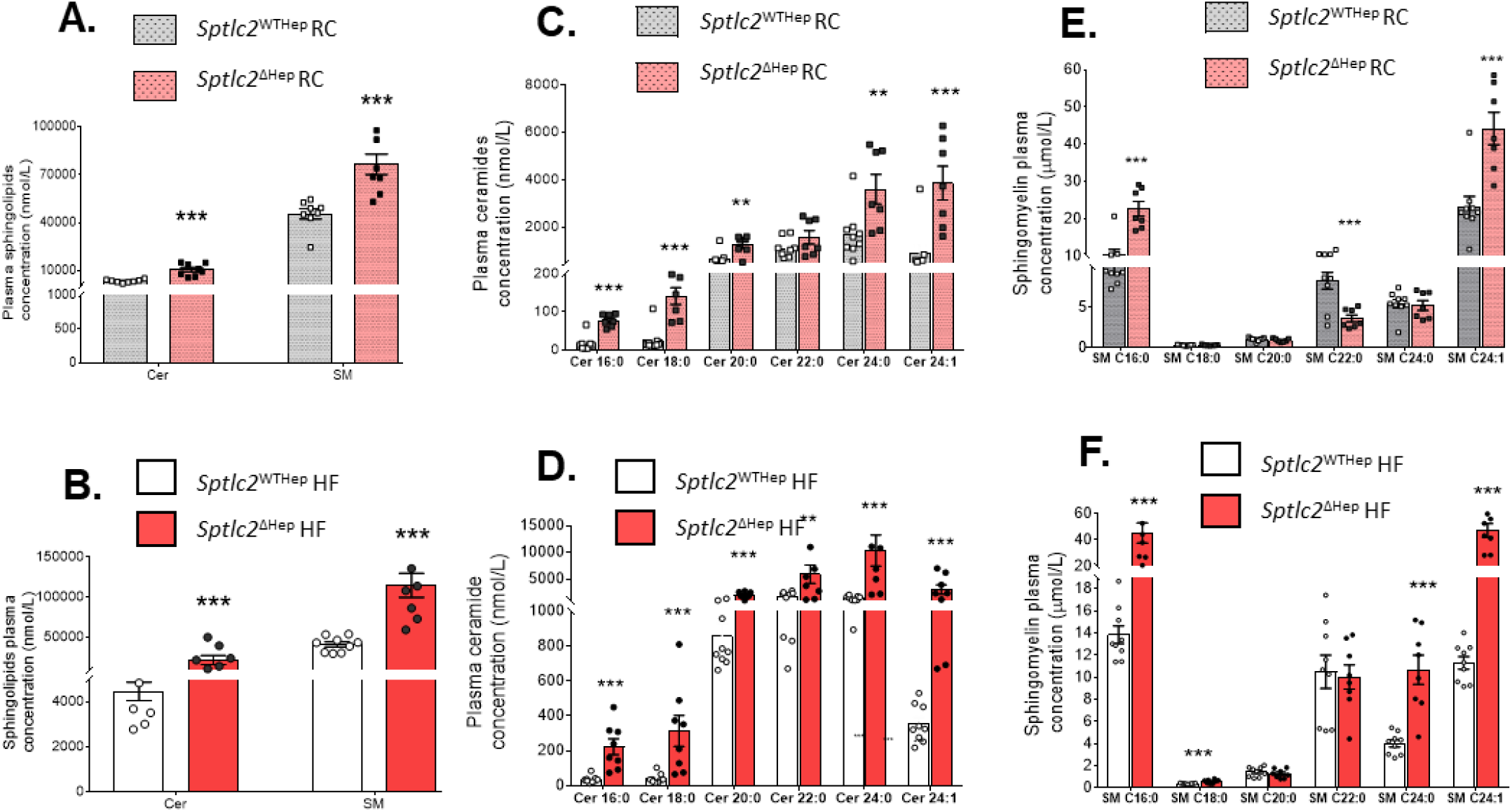
Ceramides and sphingomyelins quantification in plasma of *Sptlc2*^ΔHep^ mice and their control upon RC or HFD. (**A-B**) Plasma SM and Cer levels in regular chow (RC) mice and high fat diet (HFD) mice. (**C-D**) Plasma ceramides distribution according to acyl-chain length in RC mice and HFD mice. (**E-F**) Plasma SM distribution according to acyl-chain length in RC mice and HFD mice. Analysis were performed in 4 month-old mice and HFD mice were fed with HFD for 2 months starting at 2 month-old; error bars represents SEM; n = 6 −10 mice per groups; *p < 0.05, **p < 0.01, ***p < 0.001 via Mann-Whitney U test or Student’s *t* when appropriate. **RC**: Regular Chow; **HF**: High Fat; **Cer**: ceramide; **SM**: Sphingomyelin; **Sptlc2**: Serine Palmitoyltransferase long chain 2

### 2.2 *Sptlc2* deficiency in hepatocytes impairs bile acids homeostasis leading to a defect in lipids absorption and prevent obesity

We placed *Sptlc2*^ΔHep^ mice and their littermate controls on HFD for 8 weeks in an attempt to induce obesity and insulin resistance. As shown in Figure 2.A and 2.B, starting from week 4, mutant mice upon HFD have significantly gained less body weight than control animals, despite similar food intake (Figure 2.C). Mass composition analysis revealed a decreased fat mass in KD mice compared to WT mice upon RC and HFD (Figure 2.D). Moreover, energetic density of faeces collected every 24h during 15 days is increased in KD mice (Figure 2.E), and we showed that *Sptlc2* deficiency in hepatocytes impaired triglyceride absorption. Indeed, triglycerides amount is reduced in the plasma of *Sptlc2*^ΔHep^ mice up to 6 hours after an oral olive oil charge (Figure 2.F). These results suggest that *Sptlc2*^ΔHep^ mice exhibit a defect of lipids absorption leading to resistance to body weight gain normally induced by HFD.

**Figure 2.**
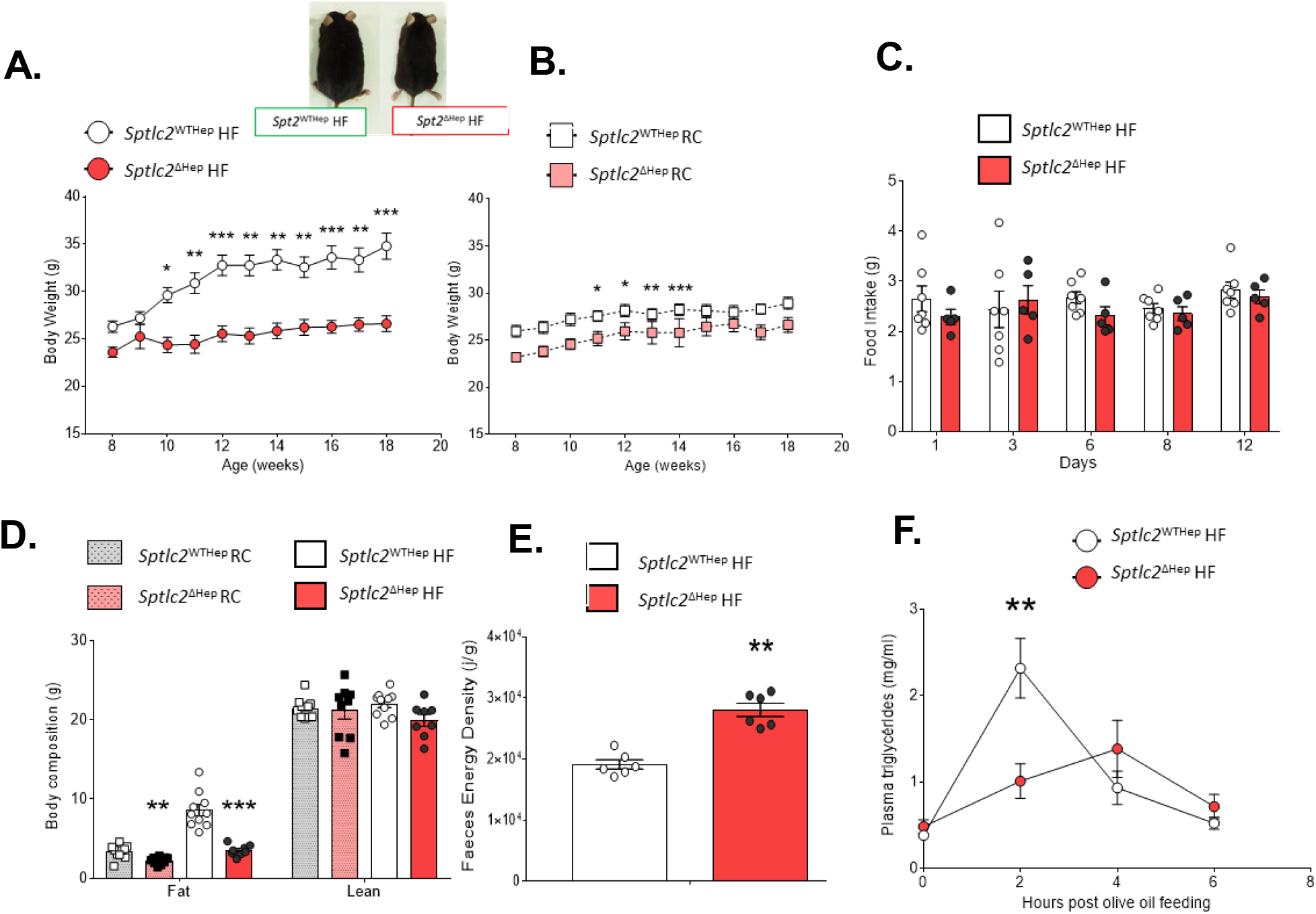
Genetic deletion of *Sptlc2* in hepatocytes prevents obesity induced by high fat diet. (**A-B**) Body weight change of *Sptlc2*^ΔHep^ and their littermate controls upon HFD or RC. (**C**) Time course of food intake of HFD mice. (**D**) Body composition, fat and lean in grams. (**E**) Energy density of faeces collected every 24H of HFD mice. (**F**) Plasma triglycerides measurement after oral olive oil charge in HFD mice. Analysis were performed in 4 months-old mice and HFD mice were fed with HFD for 2 months starting at 2 month-old; n = 4 −10 mice per groups; error bars represents SEM; *p < 0.05, **p < 0.01, ***p < 0.001 via Mann-Whitney U test, Student’s *t* test when appropriate or two-way ANOVA followed by two-by-two comparisons using Bonferroni’s post hoc test. **RC**: Regular chow; **HF**: High fat

As modifications of BA composition, synthesis and transport could impair lipid absorption (Holt 1972), we measured BA composition and concentration in the gallbladder bile, liver, ileum and plasma and showed that KD mice compared to WT mice exhibited substantially elevated BA concentrations in liver, gallbladder bile and plasma (Figure 3.A, Figure 3 – figure supplement 1.A) consistent with the reduction of bile flow (i.e. cholestasis) described in prior studies (Li et al. 2016). Moreover, we show in this study that BA composition was strongly modified between the two groups of mice, independently of the diet (Figure 3.B, Figure 3 – figure supplement 1.B). A dramatic increase in β-MCA, the dominant form of muricholic acid and in its preponderant conjugated form, tauro-muricholic acid (T-MCA) was observed in mutant mice (Figure 3.B and 3.C, Figure 3 – figure supplement 1.B and 1.C) while secondary BA, most hydrophobic BA, were almost absent. Except in the plasma, cholic acid concentration was also reduced in enterohepatic tissues in *Sptlc2*^ΔHep^ mice (Figure 3.B, Figure 3 – figure supplement 1.B). In line with these data, the hydrophobicity index of BA, was decreased in liver and gallbladder bile from *Sptlc2*^ΔHep^ mice as compared with WT mice (Figure 3.D, Figure 3 – figure supplement 1.D). Elevated plasma BA were associated with severe jaundice in mutant mice, along with moderate cytolysis and strong elevation of alkalin phosphatase and bilirubin (Figure 3.E and Table 1.), consistent with cholestasis, as reported (Li et al. 2016).

**Figure 3.**
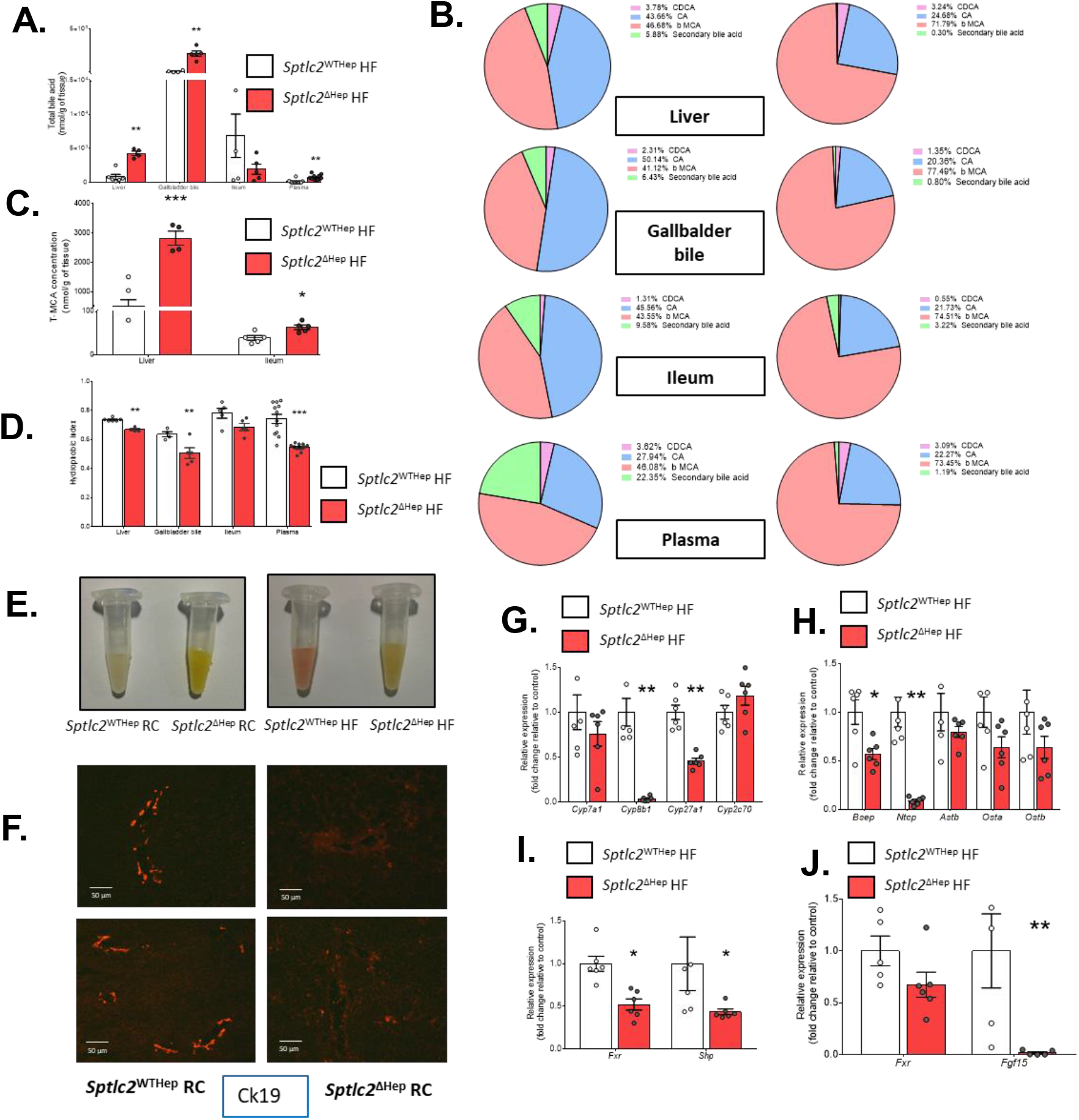
Genetic deletion of *Sptlc2* in hepatocytes impairs BA composition and metabolism. (**A**) Total bile acids content in liver, gallbladder bile, ileum and plasma of HFD mice. (**B**) Bile acids composition in gallbladder bile, ileum and plasma of HFD mutant and control mice. (**C**) Tauro-muricholic acid concentration in liver and ileum. (**D**) Hydrophobicity index of bile acids pool contained in liver, gallbladder bile, intestine and plasma of HFD *Sptlc2*^WTHep^ and *Sptlc2*^ΔHep^ mice. (**E**) Photograph of representative plasma samples from *Sptlc2*^WTHep^ or control mice upon RC or HFD. (**F**) Representative immunofluorescence image showing cytokeratin 19 (Ck19) in liver sections of RC 3 month-old mice (**G**) Relative expression of main enzymes involved in bile acids synthesis: classical or alternative pathway in liver of HFD mice, data are expressed in fold change relative to control. (**H**) Relative expression of bile acids transporters in the liver (*Bsep, Ntcp*) or terminal ileum (*Astb, Ostα/β*) of HFD mice, data are expressed in fold change relative to control. (**I-J**) Relative expression of *Fxr* and its target gene in the liver (*Shp*) or terminal ileum (*Fgf15*) of HFD mice, data are expressed in fold change relative to control. Analysis were performed in 4 months-old mice and HFD mice were fed with HFD for 2 months starting at 2 month-old; n = 4 −10 mice per groups; error bars represents SEM; *p < 0.05, **p < 0.01, ***p < 0.001 via Mann-Whitney U test, Student’s *t* test when appropriate or two-way ANOVA followed by two-by-two comparisons using Bonferroni’s post hoc test. **RC**: Regular Chow; **HF**: High Fat; CA: Cholic Acid; CDCA: Chenodeoxycholic Acid; β-MCA: beta – Murocholic Acid; **Ck19**: Cytokeratin 19; **Cyp7a1**: Cholesterol 7-alpha Hydroxylase; **Cyp8b1**: Cytochrome P450 family 8 subfamily B member 1; **Cyp27a1**: Cytochrome P450 family 27 subfamily A member 1; **Cyp2c70**: Cytochrome P450 family 2 subfamily C member 70; **Bsep**: Bile Salt Export Pump; **Ntcp**: Sodium/Taurocholate Cotransporting Polypeptide; **Asbt**: Apical Sodium-Bile acid Transporter; **Ostα/β**: Organic Solute Transporter alpha/beta; **Fxr**: Farnesoïd X Receptor; **Shp**: Small Heterodimer Partner; **Fgf15**: Fibroblast Growth Factor 15.

**Table 1.** Plasma marker of hepatic cholestasis and liver injury in *Sptlc2* ^WTHep^ HF and *Sptlc2* ^ΔHep^ HF. Analysis were performed in 4 months-old mice and HFD mice were fed with HFD for 2 months starting at 2 months-old; values are means ± SEM; n = 6 mice per groups.

|  | *Sptlc2*<sup>WTHep</sup> HF | *Sptlc2*<sup>ΔHep</sup> HF | p-value<br>(Mann-Whitney U test) |
| --- | --- | --- | --- |
| Albumin (g/L) | 28.3 ± 0.4189 | 27.3 ± 0.5795 | 0.243 |
| ALAT (U/L) | 47.6 ± 8.165 | 167.9 ± 11.99 | 0,00002 *** |
| ASAT (U/L) | 140.7 ± 26.59 | 323.5 ± 49.09 | 0.00001*** |
| Direct Bilirubin (umol/L) | 0.6 ± 0.1017 | 15.5 ± 3.929 | 0,0003 *** |
| Alkaline Phosphatase (U/L) | 42.1 ± 3.247 | 469.1 ± 57.53 | 0,0003 *** |

Interestingly in this cholestatic context, mutant mice exhibited disrupted localization of CK19 immunostaining (a marker of cholangiocytes) (Figure 3.F), suggesting that cholangiocyte injury occurred in these mice and may contribute to bile flow impairment, i.e. to cholestasis. Furthermore, we measured mRNA levels of BA transporters and the main enzymes involved in BA synthesis. Expression level of *Cyp8ab1*, which allows CA synthesis, and *Cyp27a1*, involved in the alternative pathway were both reduced in *Sptlc2*^ΔHep^ mice (Figure 3.G). Prior study has reported alterations in BA transporters localisation and hepatocyte polarity in liver of hepatocyte-specific Sptlc2 KO mice (Li et al. 2016). Here, we showed that mRNA level of the BA transporter *Bsep*, which allows BA excretion from hepatocytes to bile canaliculus and the transporter *Ntcp*, a sodium-dependent transporter expressed on the basolateral membrane of hepatocytes, responsible for BA uptake from sinusoids, are both decreased in the liver of KD mice (Figure 3.H). The significant drop of *Ntcp* expression could be part of an adaptive response to BA accumulation in hepatocytes as already well described (Zollner et al. 2001). Consistent with the dramatic increase in T-MCA, a BA reported as an FXR antagonist (Sayin et al. 2013), a significantly decreased expression in the FXR target genes *Shp* and *Fgf15* was observed in ileum and liver from *Sptlc2*^ΔHep^ as compared with WT mice (Figure 3.I and 3.J).

Thus, *Sptlc2* deficiency in hepatocytes leads to profound changes in BA synthesis and transport associated with a defect in lipids absorption and a decreased FXR target genes expression in both ileum and liver.

**Figure 3 -figure supplement 1.**
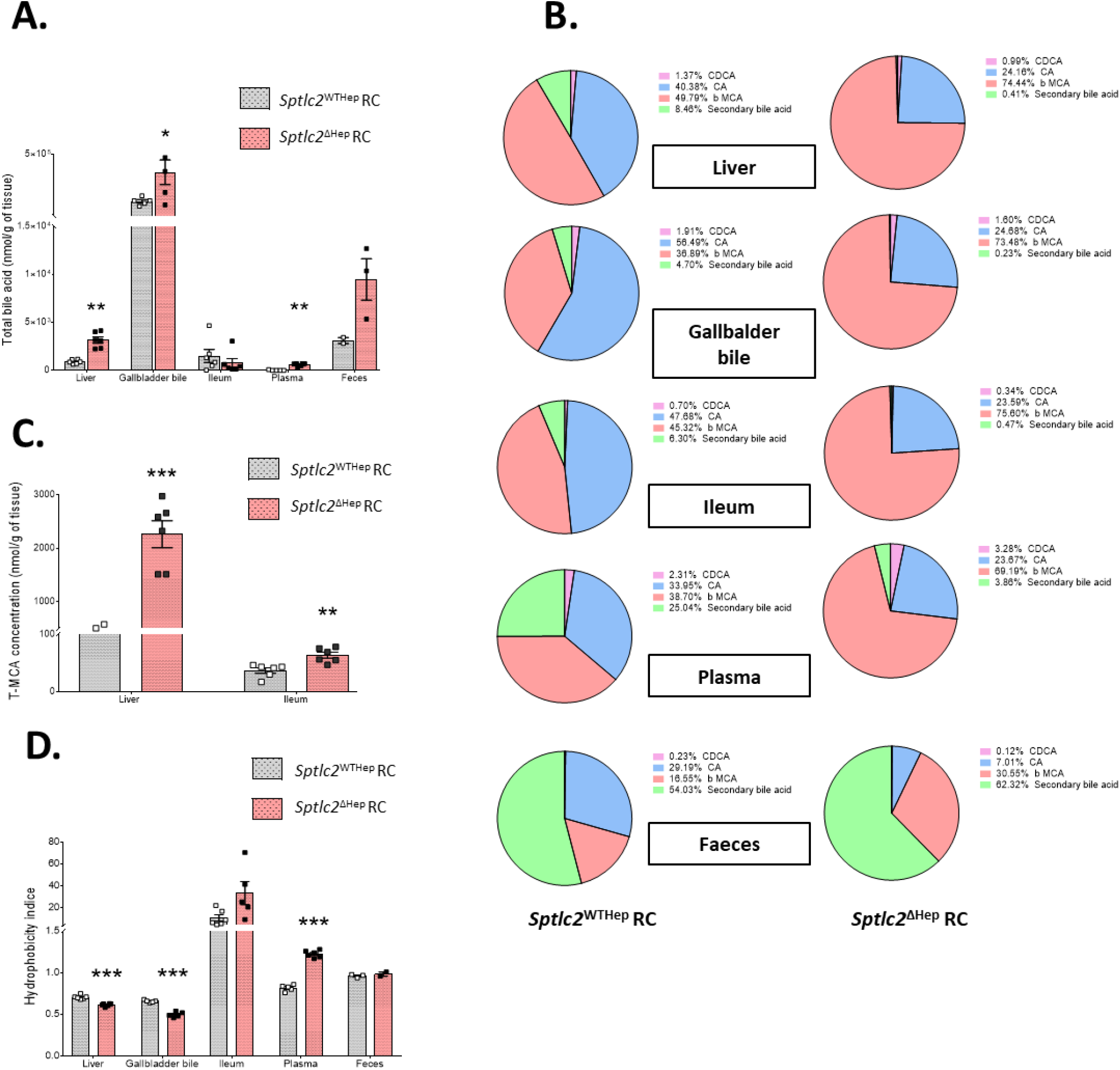
Composition of bile acids pool is altered in *Sptlc2*^ΔHep^ mice fed with control diet. (**A**) Total bile acids content in gallbladder bile, intestine and plasma of RC mice (4 month-old). (**B**) Bile acids composition in gallbladder bile, liver, ileum, plasma and faeces of RC mutant and control mice. (**C**) Tauro-muricholic acid concentration in liver and iileum. (**D**) Hydrophobicity index of bile pool contained in liver, gallbladder bile, intestine, plasma and faeces of RC *Sptlc2*^WTHep^ and *Sptlc2*^ΔHep^ mice. Analysis were performed in 4 month-old mice; error bars represents SEM; n = 3 −10 mice per groups; *p < 0.05, **p < 0.01, ***p < 0.001 via Mann-Whitney U test or Student’s *t* when appropriate. **R**C: Regular Chow; **HF:** High Fat; **CA:** Cholic Acid; **CDCA**: Chenodeoxycholic Acid; **β-MCA**: beta – Murocholic Acid.

### 2.3 Genetic deletion of *Sptlc2* in hepatocytes impairs hepatic glucose production and storage in an insulin-independent manner

The liver is at the core of glucose metabolism: it produces glucose from the breakdown of glycogen, or through gluconeogenesis from lactate, pyruvate, glycerol and amino acids (Rui 2014). These pathways are essential for normoglycemia maintenance. Interestingly, decreased level of hepatic *Sptlc2* prevents glucose intolerance in mice challenged with HFD (Figure 4.A, 4.B and 4.C). *De novo* glucose production from pyruvate (Figure 4.D, 4.E and 4.F) and glycerol (Figure 4.G, 4.H and 4.I) are decreased both in KD mice fed with HFD or RC. This observed defect in hepatic glucose production is supported by glucose production measurements on isolated hepatocytes (Figure 4.J). Two and three hours after hepatocyte stimulation induced by medium without glucose containing pyruvate and glycerol, we detected a tendency of a reduced glucose production in isolated hepatocytes from *Sptlc2*^ΔHep^ mice with or without cAMP stimulation. Indeed, hepatic *Sptlc2* deficiency decreases *G6pase* and *Pc* expression levels, two enzymes involved in glycogenesis and/or gluconeogenesis, while *Pepck* expression is unchanged (Figure 4.K). Furthermore, G6Pase activity and glucose-6-phosphate (G6P) content were decreased in mutant mice liver (Figure 4.L and 4.M). Reduced liver glycogen content was also observed in *Sptlc2* ^ΔHep^ mice at fed state or at 5h fasted state, as compared with WT mice (Figure 4.N and 4.O). Thus, consistent with these results, 6-hours-fasting glycemia is reduced in KD mice (Figure 4 – figure supplement 1).

**Figure 4.**
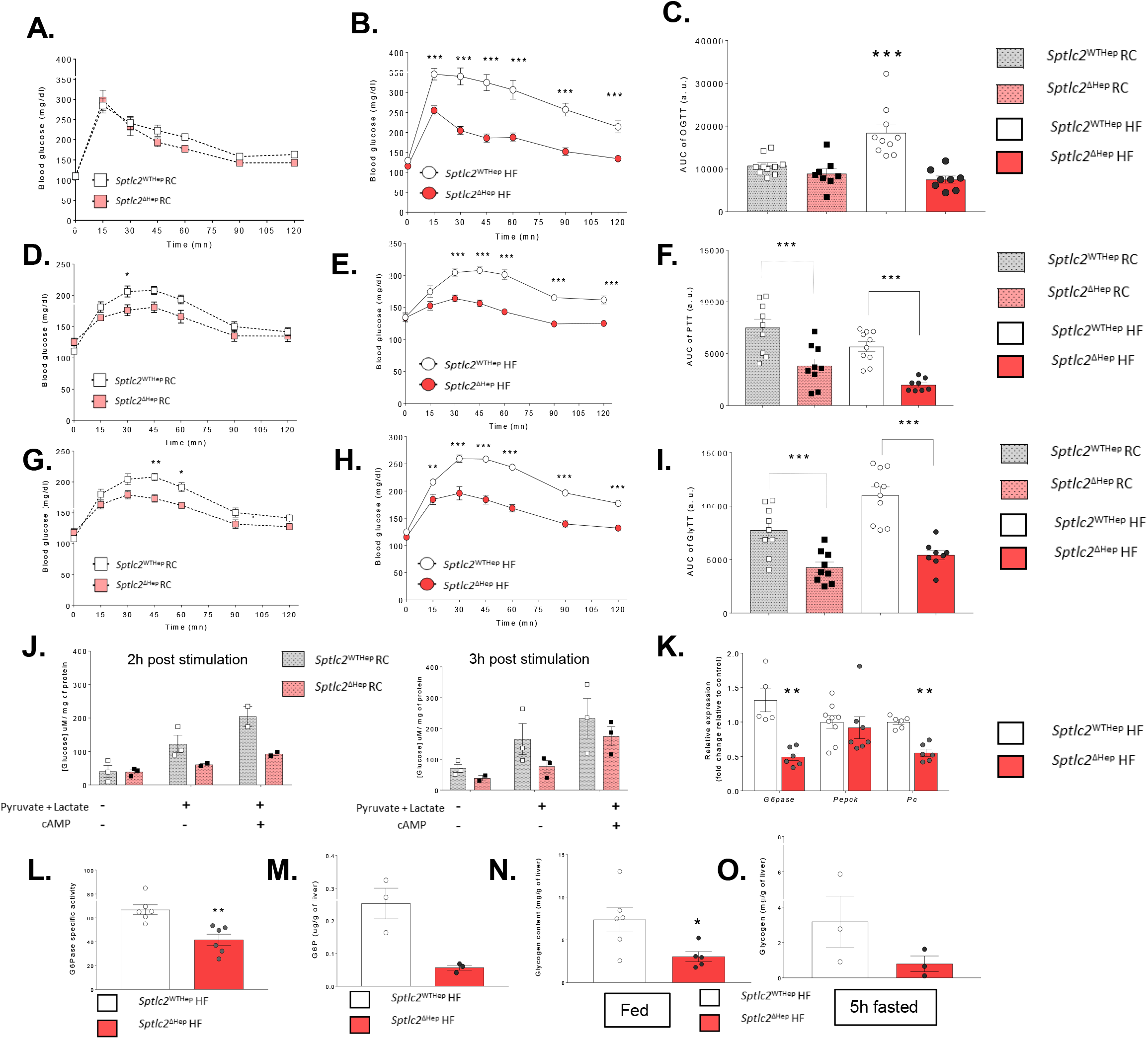
Genetic deletion of *Sptlc2* in hepatocytes impairs hepatic glucose production and storage. (**A-B**) Oral glucose tolerance test (OGTT) at 2g/kg performed in RC or HFD overnight fasting mice. (**C**) Respective areas under the curve for the OGTT in RC or HFD mice. (**D-E**) Pyruvate tolerance test (PTT) at 2g/kg performed in RC or HFD overnight fasting mice. (**F**) Respective areas under the curve for the PTT in RC or HFD mice. (**G-H**) Glycerol tolerance test (GlyTT) at 1g/kg performed in RC or HFD overnight fasting mice. (**I**) Respective areas under the curve for the GlyTT in RC or HFD mice. (**J**) Glucose concentration measured in primary hepatocytes medium after stimulation with pyruvate, lactate and cAMP (as a positive control). Primary hepatocytes were isolated from 45-day-old RC mice. (**K**) Relative expression of main enzymes involved in gluconeogenesis in liver of HFD mice, data are expressed in fold change relative to control. (**L-M**) G6Pase specific activity and G6P content in liver of HFD mice. **(N-O**) Glycogen content in liver of HFD fed or 5-hours-fasting mice Analysis were performed in 45 day-old mice (for primary hepatocytes collection) or 4 month-old mice and HFD mice were fed with HFD for 2 months starting at 2 month-old; n = 3 −10 mice per groups; error bars represents SEM; *p < 0.05, **p < 0.01, ***p < 0.001 via Mann-Whitney U test, Student’s *t* when appropriate or two-way ANOVA followed by two-by-two comparisons using Bonferroni’s post hoc test. **RC**: Regular Chow; **HF**: High Fat; **cAMP**: cyclic Adenosine MonoPhosphate; **G6Pase**: Glucose-6-Phosphatase; **Pepck**: Phosphoenolpyruvate carboxykinase; **Pc**: Pyruvate Carboxylase; **G6P**: Glucose-6-Phosphate.

**Figure 4 - figure supplement 1.**
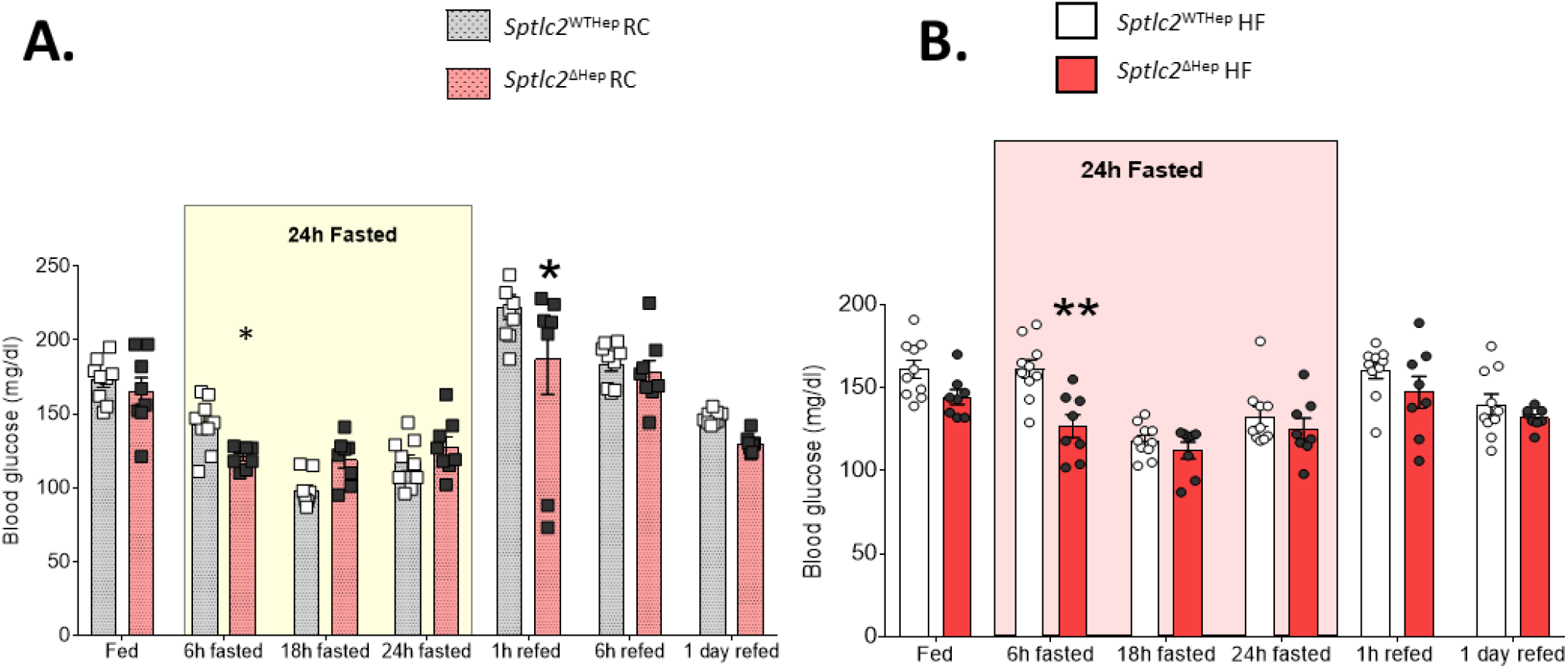
Genetic deletion of *Sptlc2* in hepatocytes led to glycemia reduction in 6-hours fasting mice. (**A-B**) Time course of glycemia measurement during a 24h fast performed in RC or HFD mice. Analysis were performed in 4 month-old mice and HFD mice were fed with HFD for 2 months starting at 2 month-old; error bars represents SEM; n = 3 −10 mice per groups; *p < 0.05, **p < 0.01, ***p < 0.001 via Mann-Whitney U test, Student’s *t* when appropriate or two-way ANOVA followed by two-by-two comparisons using Bonferroni’s post hoc test. **RC**: Regular Chow; **HF**: High Fat

In order to evaluate the role of insulin on hepatic glucose production in our model lacking *Sptlc2* expression in hepatocytes, we measured insulin secretion, during the course of a glucose tolerance test, and insulin sensitivity. Hepatic deletion of *Sptlc2* did not impair insulin secretion or systemic insulin sensitivity (Figure 5.A, 5.B and 5.C). Moreover, as expected based on C16:0-Cer and C18:0-Cer accumulation in the liver, activation by phosphorylation on serine 473 of Akt/PKB (*Protein Kinase B*), a main mediator of insulin’s anabolic effects is reduced in the liver of mutant mice (Figure 5.D, 5.E and 5.F). Thus, systemic insulin sensitivity is likely compensated by an enhanced Akt/PKB phosphorylation in the muscle (Extensor Digitorum Longus, EDL). Altogether, these results demonstrated that hepatic *Sptlc2* deficiency impairs hepatic glucose production in an insulinin-dependent manner.

**Figure 5.**
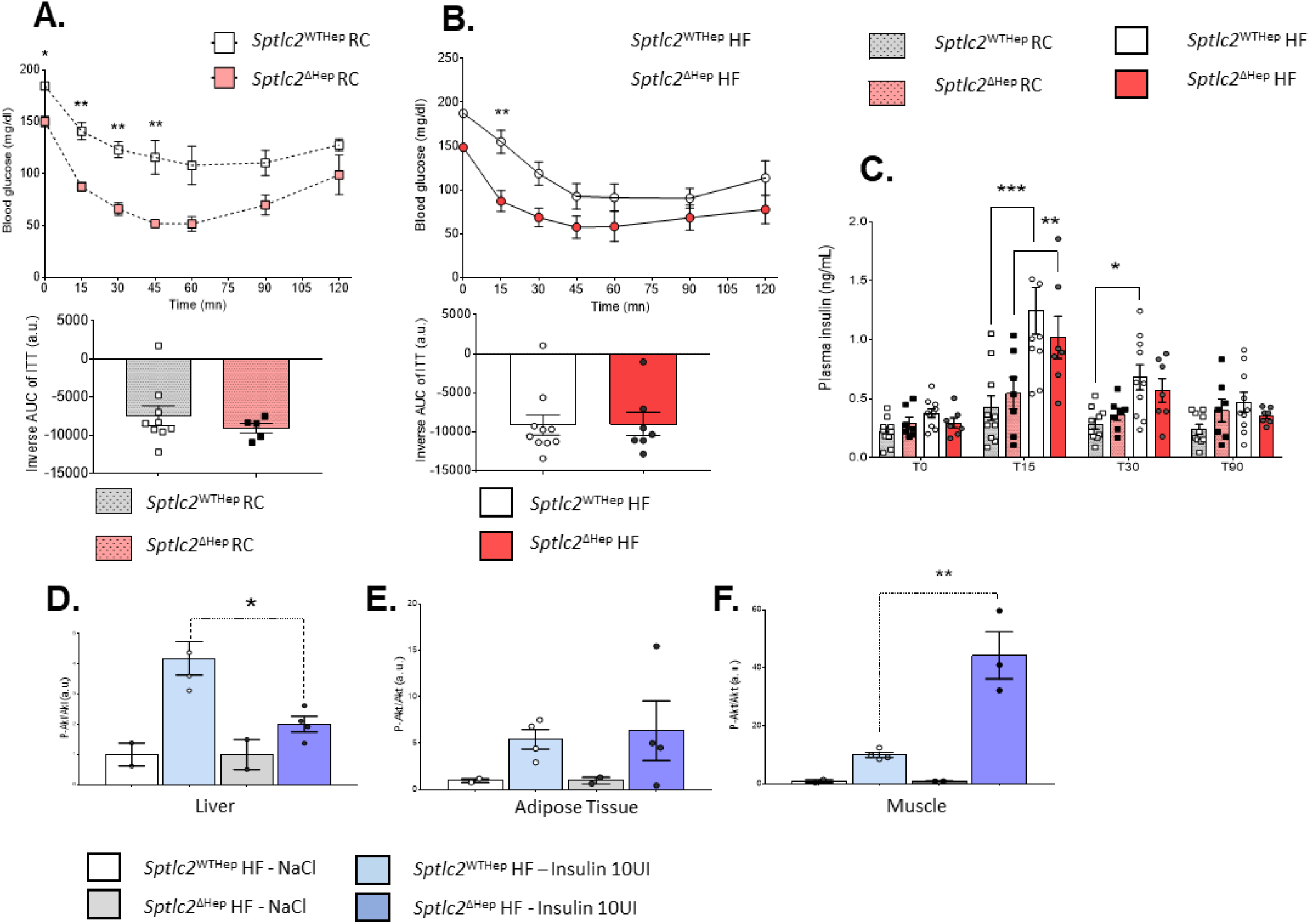
Impact of hepatic deletion of *Sptlc2* on insulin sensitivity. (**A**) Insulin tolerance test (ITT) at 0.25 UI/kg and inverse areas under the curve performed in RC mice. (**B**) Insulin tolerance test (ITT) at 0.5 UI/kg and inverse areas under the curve performed in HFD mice. (**C**) Plasma insulin secretion during OGTT test of RC or HFD mice. (**D-F**) Ratio phosphorylated Akt (Ser473)/Akt measured in cells lysates from liver, adipose tissue or muscle (extensor digitorum longus) of HFD mice stimulated with insulin (10 UI/kg) 30 mn before tissues collection. Data are expressed relative to insulin response. Analysis were performed in 45 day-old mice (for primary hepatocytes collection) or 4 month-old mice and HFD mice were fed with HFD for 2 months starting at 2 month-old; n = 3 −10 mice per groups; error bars represents SEM; *p < 0.05, **p < 0.01, ***p < 0.001 via Mann-Whitney U test, Student’s *t* when appropriate or two-way ANOVA followed by two-by-two comparisons using Bonferroni’s post hoc test. **RC**: Regular Chow; **HF**: High Fat

### 2.4 Hepatic *Sptlc2* deficiency leads to liver fibrosis, inflammation and apoptosis

Based on our data, we next asked whether modulating Cer levels could lead to liver damage. Sirius red and H&E liver section staining revealed that *Sptlc2*^ΔHep^ mice exhibited necrosis, inflammatory infiltration and fibrosis, increasing progressively with age (Figure 6.A). At 20 days, bile infarcts (i.e. necrotic cell clusters characteristic of BA overload), were observed in livers from *Sptlc2*^ΔHep^ mice but not in WT, without any significant fibrosis. At 45 days and even more at 120 days, patent fibrosis was observed, with both perisinusoidal and porto-portal accumulation of extracellular matrix (ECM). Interestingly, high fat diet doesn’t appear to exacerbate ECM accumulation in liver demonstrating strong impact of *Sptlc2* disruption on the setting of liver fibrosis. These results were associated with an increased liver weight (compared to body weight), without triglyceride accumulation, as compared to the littermate controls (Figure 6.B and 6.C).

**Figure 6.**
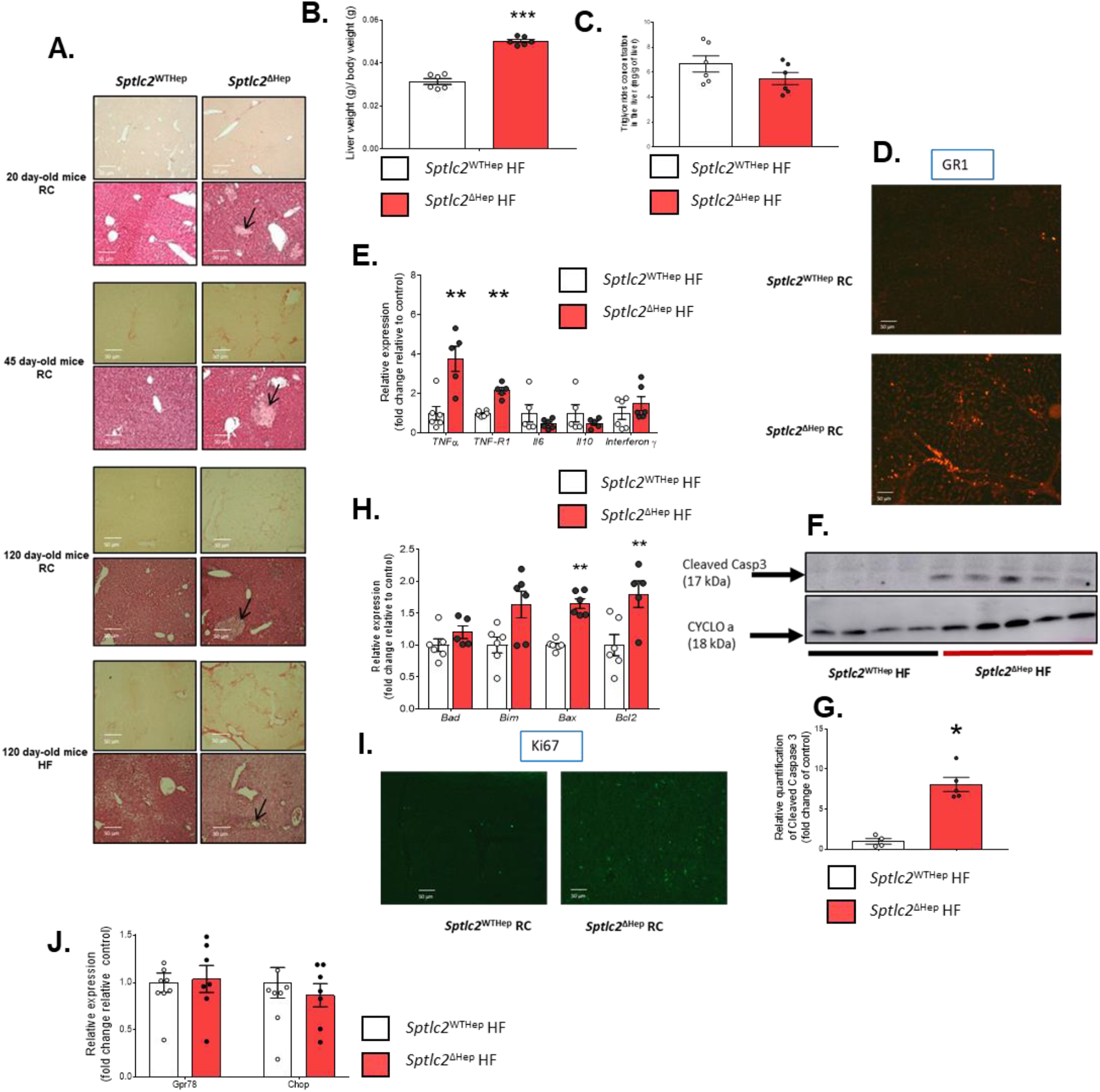
Measurement of biological parameters associated with liver damage, liver fibrosis, inflammation and apoptosis. (**A**) Representative images of H&E and sirius red staining. Images were taken in liver sections from 20 day-old RC mice, 45 day-old RC mice, 4 month-old RC mice and 4 month-old HFD mice. Black arrows represent bile infarcts. (**B**) Ratio liver weight to body weight. (**C**) Tryglicerides content in liver of HFD mice. (**D**) Representative images from immunohistochemical staining for Ly6g/GR1 in liver sections from RC mice (3 month-old). (**F-G**) Western blot analyses and quantification of cleaved caspase 3 and CYCLOa in total protein extracted from liver. (**H**) mRNA levels of *Bad, Bim, Bax* and *Bcl-2* in liver, data are expressed in fold change relative to control. (**I**) Representative images from immunohistochemical staining for the marker of proliferation, Ki67, in liver sections from RC mice (3 month-old). Expect for the immunhistochemical staining, analysis were performed in 4 month-old mice and HFD mice were fed with HFD for 2 months starting at 2 month-old; n = 4 −10 mice per groups; error bars represents SEM; *p < 0.05, **p < 0.01, ***p < 0.001 via Mann-Whitney U test, Student’s *t* when appropriate or two-way ANOVA followed by two-by-two comparisons using Bonferroni’s post hoc test. **HF**: High Fat; **Ly6g**: Lymphocyte antigen 6 complex locus G6D; **TNFα**: Tumor Necrosis Factor alpha; **TNF-R1**: Tumor Necrosis Factor Receptor 1; **Il-6**: Interleukin 6; **Il-10**: Interleukin 10; **Bad**: Bcl-2-associated death promoter; **Bim**: Bcl-2-like protein 11; **Bax**: Bcl-2-associated X protein; **Bcl-2**: β-cell lymphoma 2.

As inflammation, which could be triggered by Cer, is known to be associated with liver disorders (Seki et Schwabe 2015; Koyama et Brenner 2017), we assessed inflammatory response activation through the presence of granulocyte receptor-1 (*Gr-1*) positive cells in frozen liver tissue sections. In line with H&E staining, as shown in Figure 6.D, inflammatory *Gr1* positive cells were strongly increased in KD mice liver sections as compared with WT mice. Also, hepatic mRNAs encoded by tumor necrosis alpha (*TNFα*)-a proinflammatory cytokine also involved in apoptosis- and its TNF receptor 1 (*TNFR-1*), were significantly elevated in mice lacking expression of *Sptlc2* in the liver (Figure 6.E).

Western blot analysis revealed increased cleaved Casp3 content in the liver from HFD-fed *Sptlc2*^ΔHep^ mice as compared with WT mice (Figure 6.F and 6.G). Moreover, in *Sptlc2*^ΔHep^ mice, genes belonging to B-cell lymphoma 2 (*Bcl2*) family (apoptosis regulators), were significantly modulated. *Bcl-2* and *Bcl-2-*associated X protein (*Bax*), respectively anti-apoptotic and pro-apoptotic molecules, were up-regulated, indicating apoptosis deregulation in KD mice upon HFD (Figure 6.H). In agreement with these results, Ki67 immunostaining (a proliferation marker), revealed a dramatic increase of cell proliferation in mutant as compared with WT mice, demonstrating a high apoptosis/proliferation rate in the liver of mice lacking SPTLC2 (Figure 6.I). Interestingly, this *Sptlc2*^ΔHep^ phenotype was not associated with any ER stress, since *Gpr78* and *Chop* expression in liver were not affected by *Sptlc2* deficiency (Figure 6.J).

Longitudinal studies revealed in 20 day-old *Sptlc2*^ΔHep^ mice the presence of both bile infarcts on liver sections and altered BA pool composition (i.e increase β-MCA and decrease secondary BA), without any significant hepatic fibrosis (Figure 6 – figure supplement 1.A). In the light of these results, we can hypothesize that alteration of BA homeostasis and cholestatic disorders precede the development of hepatic fibrosis. Likewise, we observed a defect in hepatic glucose production in *Sptlc2*^ΔHep^ mice from 20 days, indicating that gluconeogenesis alteration was not related to hepatic fibrosis development (Figure 6 – figure supplement 1.B).

Altogether, these results showed that hepatic *Sptlc2* disruption led to early inflammatory liver injury, with progressive development of liver fibrosis with age, in association with abnormally high rates of apoptosis and proliferation.

**Figure 6 - figure supplement 1.**
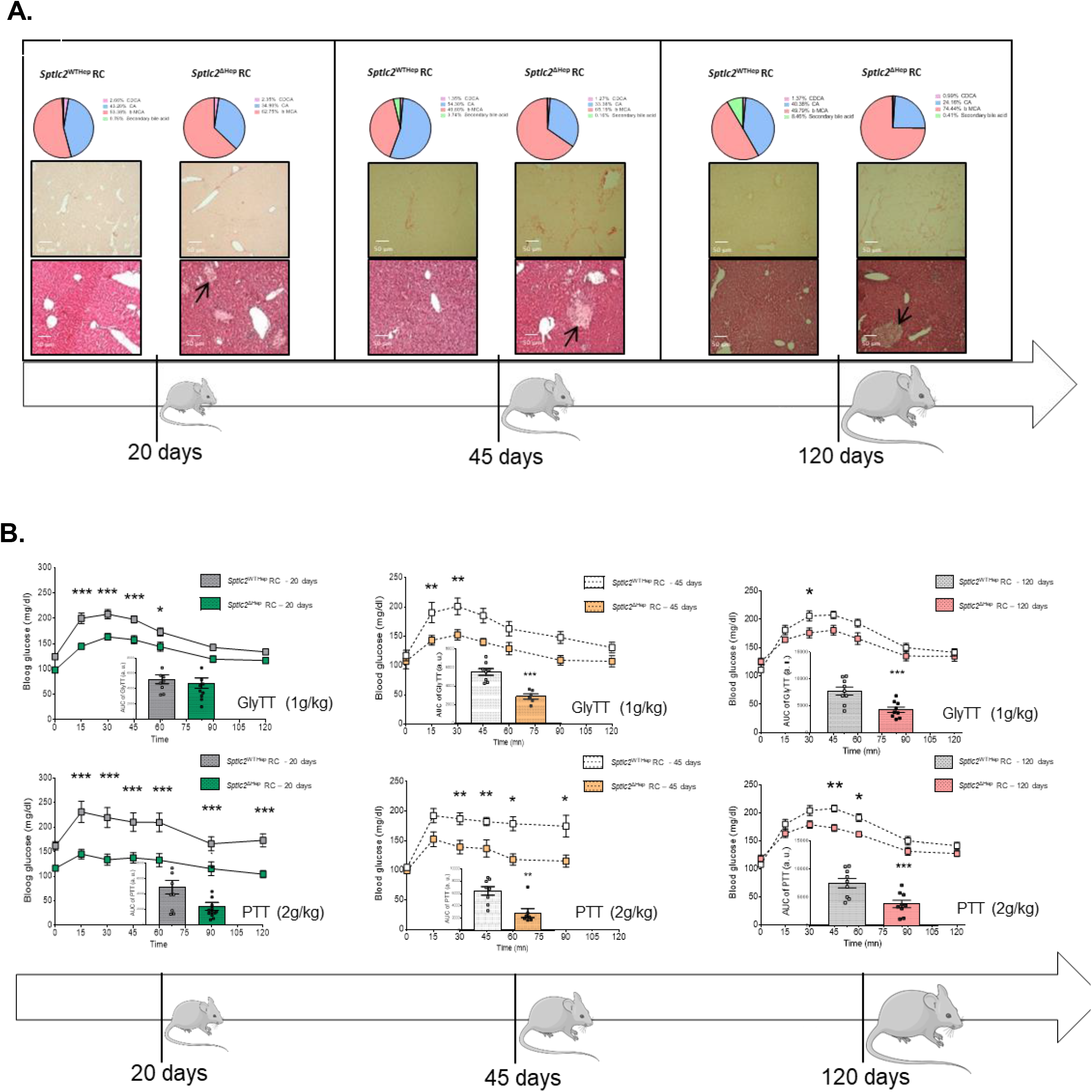
Alteration of bile acids composition and hepatic glucose production are not related to age in *Sptlc2*^ΔHep^ mice. (**A**) Progression of liver fibrosis related to age (20 day-old, 45 day-old and 120 day old) and bile acids pool composition in RC mice. Modification of bile acids pool composition and bile infarcts (represented by black arrows) are already present in livers sections from 20 day-old *Sptlc2*^ΔHep^ mice preceding the onset of fibrosis. (**B**) Evolution of hepatic glucose production from pyruvate and glycerol related to age (20 day-old, 45 day-old and 120 day-old) in RC *Sptlc2*^ΔHep^ mice and their littermate controls. **RC**: Regular Chow; **CA**: Cholic acid; **CDCA**: Chenodeoxycholic acid; **β-MCA**: beta – murocholic acid; **GlyTT**: Glycerol tolerance test; **PTT**: Pyruvate tolerance test

**Figure 6 - supplement figure 2.**
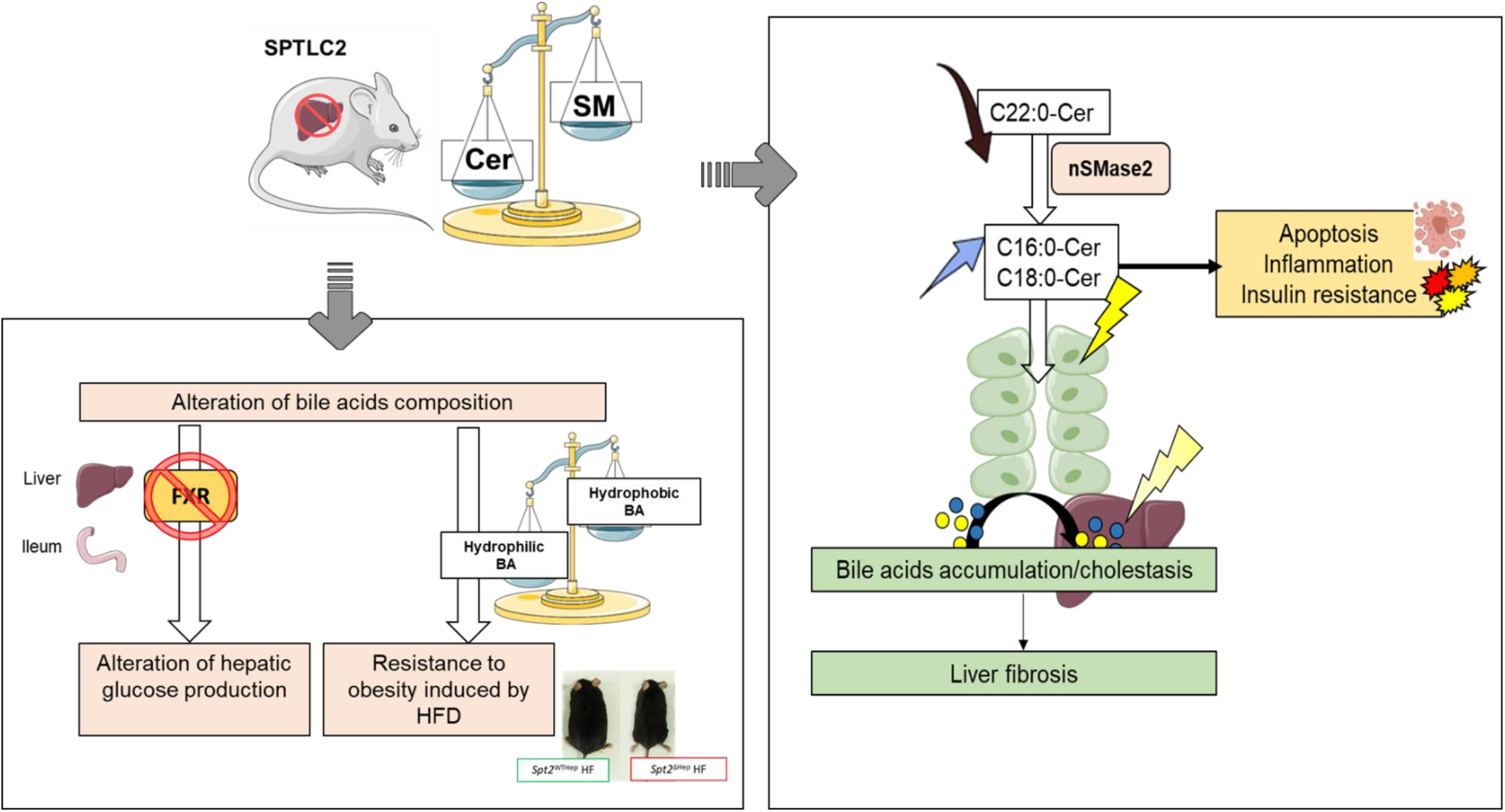
Summary of the main results from characterization of hepatocyte-specific Sptlc2 deficient mice. Genetic deletion of *Sptlc2* in hepatocytes impairs sphingomyelin/ceramide ratio in liver and induces C22:0-Cer decrease, C16:0-Cer and C18:0 increase likely through nSMase2 up-regulation. C16:0-Cer and C18:0-Cer have been described as pro-apoptosis, pro-inflammatory and inducers of insulin resistance and may cause both hepatic insulin resistance and development of liver fibrosis in *Sptlc2*^ΔHep^ mice. Hepatocyte-specific *Sptlc2* disruption alters composition of BA pool in favour of hydrophilic BA, leading to resistance to obesity linked to HFD, and in favour of T-MCA, antagonist of FXR pathway. *Sptlc2*^ΔHep^ mice exhibit a defect of hepatic glucose production, which is supposedly induced by inhibition of FXR pathway in ileum and liver. **Cer**: Ceramide; **SM**: Sphingomyelin; **Sptlc2:** Serine Palmitoyltransferase 2; **BA**: Bile Acid; **FXR**: Farnesoïd X Receptor; **HF**: High Fat

## 3. Discussion

We demonstrate for the first time a potential compensatory mechanism to regulate hepatic ceramides (Cer) content from sphingomyelin (SM) hydrolysis, and highlight the role of hepatic sphingolipid (SL) modulation on the setting of hepatic fibrosis, bile acids (BA) composition changes and hepatic glucose production (Figure 6 – supplement figure 2)

We investigated the role of Cer *de novo* synthesis in liver on energy homeostasis in mice upon regular chow (RC) or high fat diet (HFD). Interestingly, genetic deletion in hepatocytes of the rate-limiting enzyme of Cer *de novo* synthesis, *Sptlc2*, does not reduce total intrahepatic and plasma Cer level, and even increases liver Cer content in HFD fed mice. However, we show a significant decrease of the C22:0-Cer specie in liver, and both C22:0-Cer and C24:0-Cer diminution is evidenced in primary hepatocytes. These Cer species are the most abundant in liver, partially because of the high expression level of *CerS2* involved in C22:0-Cer and C24:0-Cer formation from dihydrosphingosine (Laviad et al. 2008). Thus, in our model, *Sptlc2* deficiency reduces Cer production from *de novo* synthesis for the benefit of another production pathway. Indeed, neutral sphingomyelinase 2 (nSMase2) mRNA and protein levels are up-regulated in *Sptlc2*^ΔHep^ mice, suggesting over-activation of the sphingomyelinase pathway at plasma membrane. Our data suggest for the first time to our knowledge a compensatory mechanism to maintain (and even to increase) Cer production though the dramatic up-regulation of *nSmase2* expression.

As a direct regulatory interaction between SPTLC2 and nSMase2 is unlikely due to different intracellular localization, one could speculate that the decreased utilization of serine and palmitoyl-CoA (SPTLC2 substrates) led to a modulation of *nSmase2* transcription and synthesis. Although the fine mechanisms leading to an overproduction of Cer remain unclear, these data suggest that Cer are essential components for the cell and that an attempt to alter their *de novo* synthesis would trigger a cellular response to avoid a drop in Cer content, at the expense of SM concentration, a situation occurring in liver that also leads to important changes in plasma lipids profile.

Interestingly, acute reduction of *Sptlc2* expression or activity using adenovirus targeting *Sptlc2* in liver, or pharmacological inhibitors such as myriocin, does not induce Cer accumulation (Holland et al. 2007; Li et al. 2009; Zabielski et al. 2019) in contrast to chronic *Sptlc2* deletion using genetic approach which led to Cer and especially C16:0-Cer increase in liver and adipose tissue (Li et al. 2016; S.-Y. Lee et al. 2017). In the liver, authors have shown that specific deletion of *Sptlc2* induces SM reduction and Cer accumulation in plasma membrane of hepatocytes, supporting the hypothesis of a compensatory mechanism through over-activation of SM hydrolysis at plasma membrane (Li et al. 2016). Interestingly, CerS2 null mice displays changes in the acyl chain composition of Cer, which follow the same unexpected mechanism of regulation of Cer synthesis. Indeed, liver from CerS2 null mice contains a reduced amount of C22:0-Cer and C24:0-Cer associated with an increase of C16:0-Cer leading to an unaltered total Cer content in liver of CerS2 null mice. In addition, the authors have shown an up-regulation of *Cers5* and *nSmase2* gene expression. These results corroborate the existence of a compensatory mechanism between the different pathways of Cer synthesis in liver (Pewzner-Jung, Park, et al. 2010).

In our hands, hepatic C16:0-Cer are dramatically increased in mutant mice and these Cer species are well described as apoptotic inducers and inflammation triggers (Grösch, Schiffmann, et Geisslinger 2012). Numerous studies have shown that induction of *nSmase2* expression caused C16:0-Cer accumulation, in the liver or in ileum (B. X. Wu, Clarke, et Hannun 2010; Matsubara et al. 2011). In our model, C16:0-Cer accumulation could be associated with C16:0-SM increase in plasma and, thus, resulting of SM hydrolysis, provided by the plasma, at plasma membrane in hepatocytes. This hypothesis suggest existence of two distinct pools of Cer (in plasma membrane and endoplasmic reticulum) with likely different cellular functions.

However, we observed the up-regulation of *CerS5* and *CerS6*, which allows C16:0-Cer and C18:0-Cer formation from dihydrosphingosine. In addition, neutral SMase 2 is known to show a preference for degradation of very long acyl chain SM (Marchesini et al. 2004) which are reduced in the liver and isolated hepatocytes of *Sptlc2* deficient mice. Thus, C16:0 and C18:0 accumulation could be also resulting of *CerS5* and *CerS6* up-regulation and their substrates could likely provided by SM hydrolysis and ceramidase action (He et al. 2003).

In addition, numerous studies show that *nSmase2* activation induces apoptosis through different signalling pathways leading to caspase 3 (*Casp3*) cleavage (Andrieu-Abadie et Levade 2002; Shamseddine, Airola, et Hannun 2015), consistently with our findings in *Sptlc2* ^ΔHep^ mice under HFD. Consistent with these results, CerS2 null mice display, from about 30 days of age, increased rates of hepatocyte apoptosis and proliferation leading to formation of nodules of regenerative hepatocellular hyperplasia, hepatomegaly and development of severe hepatopathy which could be associated with C16:0-Cer accumulation and C22:0-Cer reduction in liver. Interestingly, in this study, despite a total knock-out, apoptosis is restricted to the liver and, inflammatory process and fibrosis development are not observed (Pewzner-Jung, Brenner, et al. 2010).

We also showed that the modulation of sphingolipid metabolism in liver leads to a strong alteration of liver homeostasis, in particular we evidenced an impairment of BA pool composition associated with a defect in intestinal lipid absorption. BA facilitate dietary lipids uptake in the intestine and also act as signalling molecules through their action on membrane or nuclear receptors such as FXR or TGR5 (Makishima et al. 1999; Kawamata et al. 2003). Our findings show that primary BA, especially tauro-muricholic acid (T-MCA), are increased in *Sptlc2*^ΔHep^ mice and that secondary BA are almost absent. Secondary BA, which are more hydrophobic than primary BA, are essential for intestinal lipid absorption (Woollett et al. 2004). Accordingly, we measured lower hydrophobic index in BA pool of *Sptlc2* ^ΔHep^ mice leading to lower triglycerides absorption and increased energy density in faeces. Taken together, these data are consistent with a full resistance to high-fat induced obesity in *Sptlc2*^ΔHep^ mice due to altered BA-dependent lipid absorption in the intestine. Of interest, food intake is similar in *Sptlc2* ^ΔHep^ compared to the controls, supporting ingestive events rather than nervous perturbation of food behavior.

*Sptlc2* ^ΔHep^ mice also displayed jaundice as reported (Li et al. 2016). Although precise mechanisms of cholestasis development in this model remain to be fully deciphered, our data suggest that in the lack of SPTLC2 in hepatocytes, intrahepatic bile ducts were disorganized, a feature that likely contributed to bile flow impairment. How cholangiocytes would be injured in these mice remains to be elucidated, but we speculate that the accumulation of pro-inflammatory and pro-apoptosis Cer observed in *Sptlc2* ^ΔHep^ mice might contribute to biliary epithelial damage. In particular, deleterious Cer species, which are likely contained into the bile, could affect cholangiocytes (B. J. Lee et al. 2010).

Numerous actors allow the regulation of BA metabolism, among them FXR, a nuclear transcription factor also involved in glucose homeostasis. Indeed, *in vitro* or *in vivo* modulation of FXR pathway regulates gluconeogenesis in rodent. Thus, in vitro, FXR activation induces *Pepck* expression and FOXO1 activation (Stayrook et al. 2005); *in vivo*, FXR KO mice display lower *Pepck* expression and a decreased glucose production (Duran-Sandoval et al. 2005). FXR is inhibited by T-β-MCA (Sayin et al. 2013) and regulates BA synthesis (Kong et al. 2012). Consistent with the dramatic increase of T-MCA, we show that genetic deletion of hepatic *Sptlc2* inhibits expression of FXR signaling pathway. In addition, *Sptlc2* ^ΔHep^ mice have a defect in hepatic production of glucose from pyruvate and glycerol, which probably explains, associated with better insulin sensitivity in the muscle, the better glucose tolerance under HF. Of note, 6-hours-fasting glycemia is reduced in *Sptlc2*^ΔHep^ mice. These traits are unlikely linked to a liver-specific increase in insulin sensitivity, while Cer, and especially C16:0-er and C18:0-Cer, well-known inducers of insulin resistance, accumulates in liver. Consistently, glycogen content is reduced, and phosphorylation of Akt/ protein kinase B (PKB), a main mediator of insulin anabolic effects is reduced in the liver of mutant mice, while systemic insulin sensitivity seems to be compensated by an enhanced Akt/PKB phosphorylation in the muscle. Altogether, these results demonstrated that hepatic *Sptlc2* deficiency impaired hepatic glucose production in an insulin independent way.

We also evidenced a previously undescribed age-related effect of SPTLC2 deficiency that favours hepatic fibrogenesis. Although precise mechanisms remain to be investigated, the decreased amount of membrane SM in the lack of hepatic SPTLC2 (Li et al. 2016), associated with a reduction of membrane lipids rafts, may favour beta catenin translocation to the nucleus and contribute to the onset of fibrogenesis (Ge et al. 2014; Li et al. 2016). Moreover, several studies showed that Cer themselves are involved in hepatic fibrogenesis, (Simon et al. 2019; Ishay et al. 2020). In particular, it has been shown in a mouse NASH model that a decrease in hepatic Cer content (through Fgf19 treatment that modulates BA metabolism) rescued from hepatic fibrosis, triglycerides accumulation and liver injury (Zhou et al. 2017). Interestingly, in our model we demonstrated that the hepatic glucose production defect was already present in 20 day-old *Sptlc2* ^ΔHep^ mice before the neat onset of fibrosis and altogether our results suggest that hepatic defect of glucose production is independent of fibrosis development over time.

In conclusion, our findings suggest a compensatory mechanism for Cer production in liver, and highlight a link between sphingolipid production, BA metabolism, and glucose production in an insulin-independent way. The BA nuclear receptor FXR, which is involved in both BA homeostasis and energy metabolism, is likely to be at the core of a sphingolipids/BA/neoglucogenesis axis. These data suggest a new role of nSMase2. Recently, it has been shown that *nSmase2* is a target gene of FXR (Xie et al. 2017; Q. Wu et al. 2021). Although *Fxr* and *nSmase2* vary in opposite ways in our model, our findings suggest that nSMase2 may exert a negative feedback on FXR expression, probably through its products, Cer.

## 4. Materials and methods

### 4.1 Animals

All procedures were carried out in accordance with the ethical standards of French and European regulations (European Communities Council Directive, 86/609/EEC). Animal use and procedures were approved by the Ethics committee of the University of Paris and by the French Ministry of Research under the reference #2016040414129137. *Sptlc2^lox/lox^* mice (kindly given by Dr Xian-Cheng Jiang, SUNY Downstate Health Sciences University, New York, USA) were crossed with *AlbCre*^+/-^ mice (kindly given by Dr Catherine Postic, Cochin Institute, Paris, France) to generate mice lacking *Sptlc2* expression in hepatocytes *Sptlc2*^ΔHep^ or littermate controls *Sptlc2*^WTHep^. Male *Sptlc2*^WTHep^ and *Sptlc2*^ΔHep^ were housed in a controlled environment with tap water and ad libitum regular food (A04 diet, Safe, Augy, France) or high fat diet (HF260 diet, Safe, Augy, France) starting at 8 weeks of age. Mice were maintained on a 12-hour light/12-hour dark cycle and cages were enriched with tunnels. Number of mice and suffering were minimized in accordance with the 3Rs.

### 4.2 Metabolic phenotyping

#### Body weight and body mass composition

Body weight was measured weekly. Body mass composition was determined by an EchoMRI-900 (Echo Medical Systems).

#### Oral glucose tolerance test

Oral glucose tolerance tests (OGTT) were performed in overnight fasting mice. A glucose solution (2 g/kg) was administrated by oral gavage. Blood glucose was quantified from the tip of the tail vein with a glucose meter (Glucofix Lector, Menarini Diagnostics, Rungis, France). Blood samples were collected from the tail vein to assay plasma insulin with a wide-range mouse ultrasensitive insulin ELISA Kit (catalog no. 80-INSMSU-E01, Alpco, Salem, NH, USA).

#### Insulin tolerance test

To assess insulin sensitivity, insulin tolerance tests (0.5 UI/kg for HFD mice and 0,25UI/kg for RC mice) were performed in 5-hour-fasting mice. Mice were given an intraperitoneal injection of insulin (Novo Nordisk, Bagsværd, Danemark), and blood glucose was monitored following the same protocol as was used for OGTT.

#### Pyruvate and glycerol tolerance test

To evaluate hepatic glucose production, pyruvate at 2g/kg (sodium pyruvate, catalog no. P5280, Sigma-Aldrich, Saint-Louis, MO, USA) or glycerol at 1g/kg (catalog no. G5516, Sigma-Aldrich, Saint-Louis, MO, USA) were injected (intraperitoneal injection) in overnight fasting mice. Blood glucose was measured every 15 or 30 minutes during 120 minutes and quantified from the tip of the tail vein with a glucose meter (Glucofix Lector, Menarini Diagnostics, Rungis, France).

### 4.3 Energy content of faeces by bomb calorimeter

Faeces were collected every 24h during one week, dried to constant weight (0.001 g) at 60 °C, and energy content of dried faeces (kJ/g) was analyzed for each individual mouse using a bomb calorimeter (IKA C200, Staufen, Germany).

### 4.4 Liver histology

Liver samples were either frozen in nitrogen-cooled isopentane and stored at −80°C until use or fixed in 4% formaldehyde and embedded in paraffin. H&E (Haemotoxylin and Eosin) and Sirius red staining were performed following standard procedures.

#### Immunohistochemistry

Frozen Liver samples sections 10-μm thick were cut with a refrigerated cryostat, collected on slides, and stored at −80 °C until staining. Antibodies against: Ki67 (catalog no.15580, 1/500, Abcam, Cambridge, United-Kingdom), Gr-1 (catalog no. 550291, 1/50, BD Pharmingen, Franklin Lakes, NJ, USA) and the cholangiocyte marker cytokeratin-19 CK19 (TROMA-III 1/500, Developmental Studies Hybridoma Bank, University of Iowa, IA, USA) were used. Liver sections were then incubated with secondary antibodies (Alexa fluor 1/500, Molecular Probes, Eugene, OR, USA) 30 min at 37°C. Images were acquired by epifluorescence (Axioskope, Zeiss, Oberkochen, Germany).

### 4.5 Real time PCR analysis

#### RNA extraction and cDNA synthesis

Total RNA was isolated from the liver using RNeasy Mini kit (Qiagen, Hilden, Germany). The concentration of RNA samples was ascertained by measuring optical density at 260 nm. The integrity of RNA was confirmed by the detection of 18S and 28S bands after agarose-formaldehyde gel electrophoresis. The quality of RNA was verified by optical density absorption ratio OD 260nm / OD 280nm. To remove residual DNA contamination, the RNA samples were treated with DNAse RNAse-free (Qiagen, Hilden, Germany) and purified with Rneasy mini column (Qiagen, Hilden, Germany). 1 μg of total RNA from each sample was reverse transcribed with 40 U of M-MLV Reverse Transcriptase (Thermofisher Scientific, Waltham, MA, USA) using random hexamer primers.

#### Real-time PCR using SYBR-Green chemistry

Real time quantitative PCR amplification reaction were carried out in a LightCycler 480 detection system (Roche, Basel, Switzerland) using the LightCycler FastStart DNA Master plus SYBR Green I kit (catalog n° 03515869001, Roche, Basel, Switzerland). 40ng of reverse transcribed RNA was used as template for each reaction. All reactions were carried out in duplicate with no template control. The PCR conditions were: 95°C for 10 min, followed by 40 cycles of 95°C for 10 sec, 60°C for 10 sec and 72°C for 10 sec. To compare target level, relative quantification was performed using the 2^-ΔΔCt^ methods. The relative abundances of the involved genes were calculated by normalizing their level to those of Cyclophilin A (*CycloA*), 18S ribosomal RNA *(18S)* and TATA-binding protein (*Tbp*).The primers (Thermofisher Scientific, Waltham, MA, USA) were derived from mouse sequences and are listed in supplemental methods (supplement table 2.).

### 4.6 Western blot analysis

Samples of frozen mouse liver were homogenized in lysis and extraction buffer (RIPA lysis and extraction buffer, catalog n° 89900, Thermofisher Scientific, Waltham, MA, USA) containing protease inhibitors (protease inhibitor cocktails tablets, catalog n° 48047900, Roche, Basel, Switzerland). After high-speed shaking in TissueLyserII (Qiagen, Hilden, Germany) with stainless steel beads, the homogenate was centrifuged at 15 000 rpm for 15 min at 4 °C, and the supernatant was collected in a new tube. After appropriate quantitative analysis (protein quantitation kit, Interchim, Montluçon, France), equal amounts of the protein samples (20 μg of liver extracts) were resuspended in Laemmli sample buffer (catalog n°39000, Thermofisher Scientific, Waltham, MA, USA) and separated in an 4%–20% sodium dodecylsulfate (SDS) polyacrylamide gel system (Biorad, Hercules, CA, USA). After transfer, the nitrocellulose membranes were incubated with specific antibodies overnight at 4 °C and then with the secondary antibody conjugated with peroxidase for 1 h at RT. The primary antibodies used were the following: anti-cleaved Casp3 (catalog n°9661, 1/500, Cell Signaling Technology, Danvers, MA, USA), anti-Smpd3 (Merck clone 14G5.1, 1/2000, Merck, Darmstadt, Germany), and anti-Cyclophilin A (catalog n°51418, 1/2000, Cell Signaling Technology, Danvers, MA, USA). Immunoreactivity was detected with an ECL Western Blotting Analysis System (Thermofisher Scientific, Waltham, MA, USA) and acquired and analyzed using Amersham™ Image 6000 and the Image J software (LiCor Biosciences, Lincoln, NE, USA).

### 4.7 Quantification of phosphorylated (Ser473) Akt and total Akt

Protein extraction was carried out from liver, adipose tissue and muscle following the same protocol as was used for western blot analysis. Quantitative determination of phospho-Akt (Ser473) and total Akt was assessed by a Meso Scale Discovery (MSD, Kenilworth, NJ, USA) multispot electrochemiluminescence immunoassay system (Phospho(Ser473)/Total Akt Whole Cell Lysate Kit, catalog n° K15100D) according to manufacturer’s instructions.

### 4.8 Biochemical analyses

Liver triglycerides were measured using triglyceride determination kit from Sigma (catalog n°TR0100, Sigma-Aldrich, Saint-Louis, MO, USA) according to the manufacturer’s instructions. Liver glycogen content was measured using amyloglucosidase enzyme (catalog n°A7095, Sigma-Aldrich, Saint-Louis, MO, USA) assay according to Roehrig and Allred method (Roehrig et Allred 1974). Glucose was then determined using Glucose GOD-PAP kit (catalog n°87109, Biolabo, Maizy, France). Plasma concentrations of alanine amino-transferase (ALAT), aspartate amino-transferase (ASAT), direct bilirubin and alkaline phosphatase were determined using an automated Monarch device (CEFI, IFR02, Paris, France) as described previously (Viollet et al. 2003).

### 4.9 Isolation of primary mouse hepatocytes

Hepatocytes from 45 day-old *Sptlc2*^WTHep^ and *Sptlc2*^ΔHep^ mice were isolated as previously described (Besnard et al. 2016).

### 4.10 Measurement of glucose production from isolated hepatocytes

Primary mouse hepatocytes were cultured in BD BioCoat™Collagen 6-well plates (BD Pharmingen, Franklin Lakes, NJ, USA) at 1-1.5 x 10^6^ cells per well in William’s medium (catalog n°12551032, Gibco, Thermofisher Scientific, Waltham, MA, USA) supplemented with 5% FBS, penicillin (100 units/ml) and streptomycin (100 μg/ml). The day after primary hepatocytes isolation, cells were washed two times with PBS+/+ and medium was replaced with glucose-free DMEM (catalog n°D5030, Sigma-Aldrich, Saint-Louis, MO, USA) without phenol red supplemented with 2 mM L-glutamine, 15 mM HEPES (catalog n°15630-056, Gibco, Thermofisher Scientific, Waltham, MA, USA), 5% penicillin (100 units/ml) and streptomycin (100 μg/ml) and in presence or absence (negative control) of 20 mM sodium lactate (catalog n°L7022, Sigma-Aldrich, Saint-Louis, MO, USA), 1,5 mM sodium pyruvate (catalog n°11360-070, Gibco, Thermofisher Scientific, Waltham, MA, USA) and pCPT-cAMPC (catalog n°C3912, Sigma-Aldrich, Saint-Louis, MO, USA) at 100μM as positive control. Hepatocytes were then incubated at 37 °C and 150μl of cell medium was collected 2h and 3h after stimulation for glucose measurement. The glucose content of the supernatant was measured using hexokinase (catalog n° H4502, Sigma-Aldrich, Saint-Louis, MO, USA) and glucose-6-phosphatase enzyme (catalog n°G8404, Sigma-Aldrich, Saint-Louis, MO, USA).

### 4.11 G6pase activity assay and liver G6P content measurement

G6Pase activity was assayed in liver homogenates for 10 min at 30 °C at pH 7.3 in the presence of a saturating glucose-6-phosphate concentration of 20mM as described in (Mithieux et al. 2004; Rajas et al. 1999). Hepatic glucose-6-phosphatase (G6P) determinations were carried out as previously described (Penhoat et al. 2014).

### 4.12 Quantification of sphingolipids

All solvents used were LC-MS grade and purchased from Biosolve (Valkenswaard, Netherlands). Standard compounds were obtained from Sigma Aldrich (Saint-Louis, MO, USA). Individual stock solutions (2.5 mmol/L) of sphingolipids (sphingosine-1-phosphate (S1P), Cer, hexosyl ceramides (HexoCer), lactosyl ceramides (LactoCer) and SM), and exogenous S1P (d17:1), Cer (18:1/17:0) and SM (18:1/17:0) were prepared in isopropanol. A pool of reference standard solutions including S1P, 6 Cer species, 3 HexoCer species, 3 LactoCer species and 6 SM species (Supplemental Table 1) was prepared in isopropanol and then serially diluted to obtain seven standard solutions ranging between 1-1000 nmol/L for S1P, 5-5000 nmol/L for Cer, HexoCer and LactoCer, and 0.1-100 μmol/L for SM. Liver tissues samples were weighted and diluted (0.1 g/mL) in PBS before homogenization. Standard solutions, liver homogenates and plasma samples (10 μL) were then extracted with 500 μL of methanol/chloroform mixture (2:1; v:v) containing exogenous internal standards (IS) at 0.5 μmol/L, 1 μmol/L and 5 μmol/L for S1P (17:1), Cer (18:1/17:0) and SM (18:1/17:0), respectively. Samples were mixed and centrifuged for 10 min at 20 −000× g (4 °C). The supernatants were dried under a gentle stream of nitrogen. Dried samples were finally solubilized in 500 μL of methanol prior liquid chromatography-tandem mass spectrometry (LC-MS/MS) injection.

Sphingolipid concentrations were determined in plasma and liver tissue by LC-MS/MS on a Xevo^®^ Triple-Quadrupole mass spectrometer with an electrospray ionization interface equipped with an Acquity H-Class^®^ UPLC™ device (Waters Corporation, Milford, MA, USA). Samples (10 μL) were injected onto an Acquity BEH-C_18_ column (1.7 μm; 2.1 × 50 mm, Waters Corporation) held at 60 °C, and compounds were separated with a linear gradient of mobile phase B (50% acetonitrile, 50% isopropanol containing 0.1% formic acid and 10 mmol/L ammonium formate) in mobile phase A (5% acetonitrile, 95% water containing 0.1% formic acid and 10 mmol/L ammonium formate) at a flow rate of 400 μL/min. Mobile phase B was linearly increased from 40% to 99% for 4 min, kept constant for 1.5 min, returned to the initial condition over 0.5 min, and kept constant for 2 min before the next injection. Target compounds were then detected by the mass spectrometer with the electrospray interface operating in the positive ion mode (capillary voltage, 3 kV; desolvatation gas (N_2_) flow, 650 L/h; desolvatation gas temperature, 350 °C; source temperature, 120 °C). The multiple reaction monitoring mode was applied for MS/MS detection as detailed in Supplemental Table 1. Chromatographic peak area ratios between sphingolipids and IS constituted the detector responses. Standard solutions were used to plot the calibration curves for quantification. The assay linearity was expressed by the mean *R*^2^, which was greater than 0.996 for all compounds (linear regression, 1/x weighting, origin excluded). Data acquisition and analyses were performed with MassLynx and TargetLynx version 4.1 software, respectively (Waters Corporation, Milford, MA, USA).

### 4.13 Bile acids measurements

BA measurements were performed on mouse bile, plasma, liver and faeces by high-performance liquid chromatography-tandem mass spectrometry as described in (Péan et al. 2013; Bidault-Jourdainne, Merlen et al. 2020). The hydrophobicity index reflects BA hydrophobicity, taking into account the retention time (RT) of different BA on a C18 column with a gradient of methanol; the LCA has the highest retention time, the TUDCA-3S has the lowest.

### 4.14 Statistical Analysis

Data are expressed as means ± SEM. Statistical analysis was performed using Student’s *t* test, Mann-Whitney U test or two-way ANOVA followed by two-by-two comparisons using Bonferroni’s post hoc test (GraphPad Prism 6 Software, La Jolla, CA, USA). Differences were considered significant at *p* < 0.05.

### 4.15 Materials and methods - supplement

**Supplemental table 1.** Multiple reaction monitoring (MRM) transitions used for LC-MS/MS detection.

1 4.15 Materials and methods - supplement
| Sphingolipid species | Cone/collision (V) | MRM transition (m/z) |
| --- | --- | --- |
| S1P (18:1) | 28/14 | 380.2 → 264.3 |
| S1P (17:1), IS | 28/14 | 366.2 → 250.3 |
| Cer (d18:1/16:0) | 28/26 | 538.5 → 264.3 |
| Cer (d18:1/18:0) | 30/26 | 566.5 → 264.3 |
| Cer (d18:1/20:0) | 28/26 | 594.3 → 264.3 |
| Cer (d18:1/22:0) | 30/30 | 622.6 → 264.3 |
| Cer (d18:1/24:1) | 28/30 | 648.6 → 264.3 |
| Cer (d18:1/24:0) | 34/26 | 650.6 → 264.3 |
| HexoCer (d18:1/16:0) | 26/38 | 700.5 → 264.3 |
| HexoCer (d18:1/24:1) | 26/40 | 810.7 → 264.3 |
| HexoCer (d18:1/24:0) | 26/40 | 812.8 → 264.3 |
| LactoCer (d18:1/16:0) | 30/44 | 862.6 → 264.3 |
| LactoCer (d18:1/24:1) | 32/50 | 972.7 → 264.3 |
| LactoCer (d18:1/24:0) | 36/50 | 974.7 → 264.3 |
| Cer (d18:1/17:0), IS | 28/28 | 552.5 → 264.3 |
| SM (d18:1/16:0) | 58/26 | 703.6 → 184.1 |
| SM (d18:1/18:0) | 58/32 | 731.5 → 184.1 |
| SM (d18:1/20:0) | 46/26 | 759.7 → 184.1 |
| SM (d18:1/22:0) | 40/30 | 787.7 → 184.1 |
| SM (d18:1/24:1) | 38/30 | 813.7 → 184.1 |
| SM (d18:1/24:0) | 36/30 | 815.7 → 184.1 |
| SM (d18:1/17:0), IS | 56/30 | 717.6 → 184.1 |

*IS, internal standard*

**Supplemental table 2.**
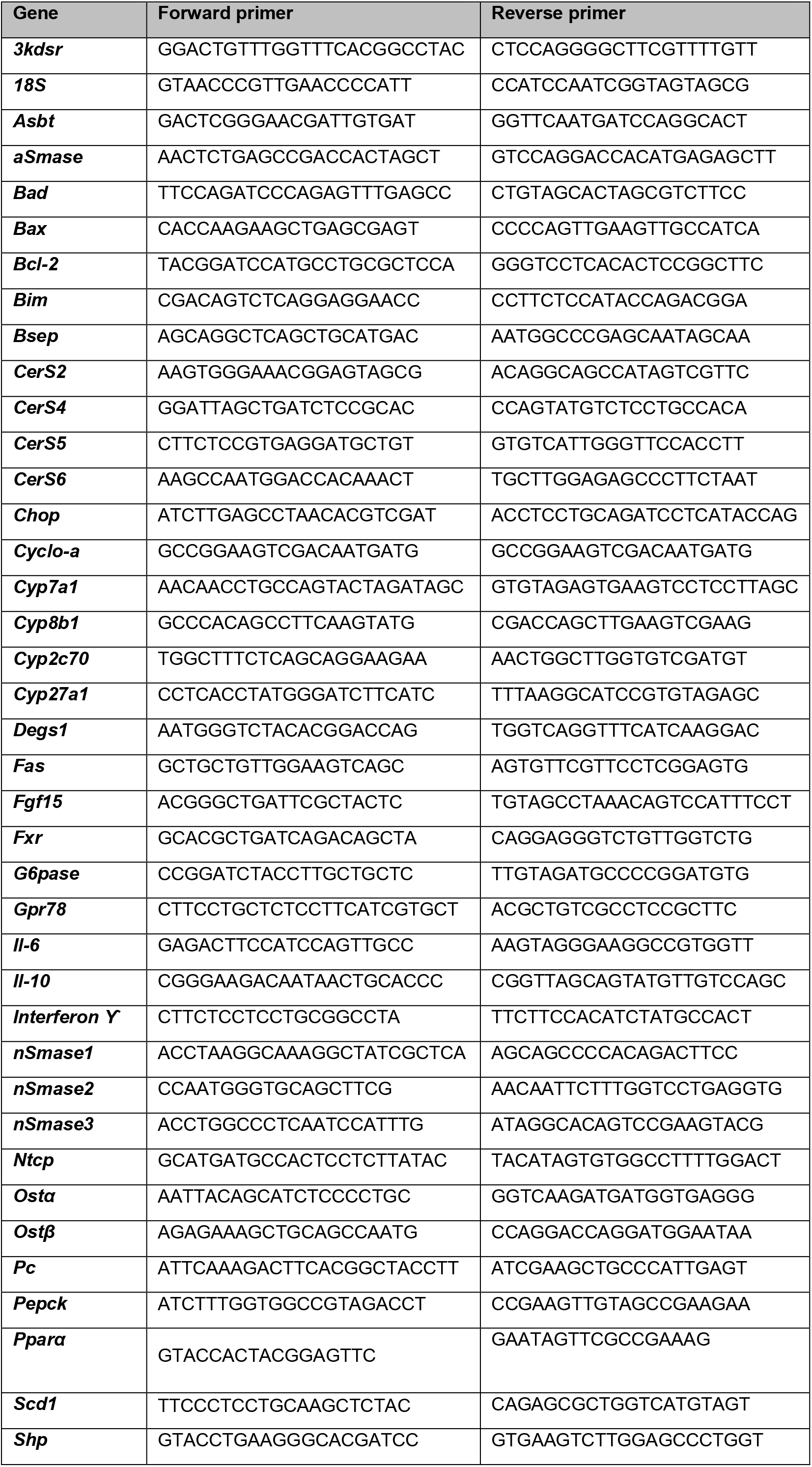
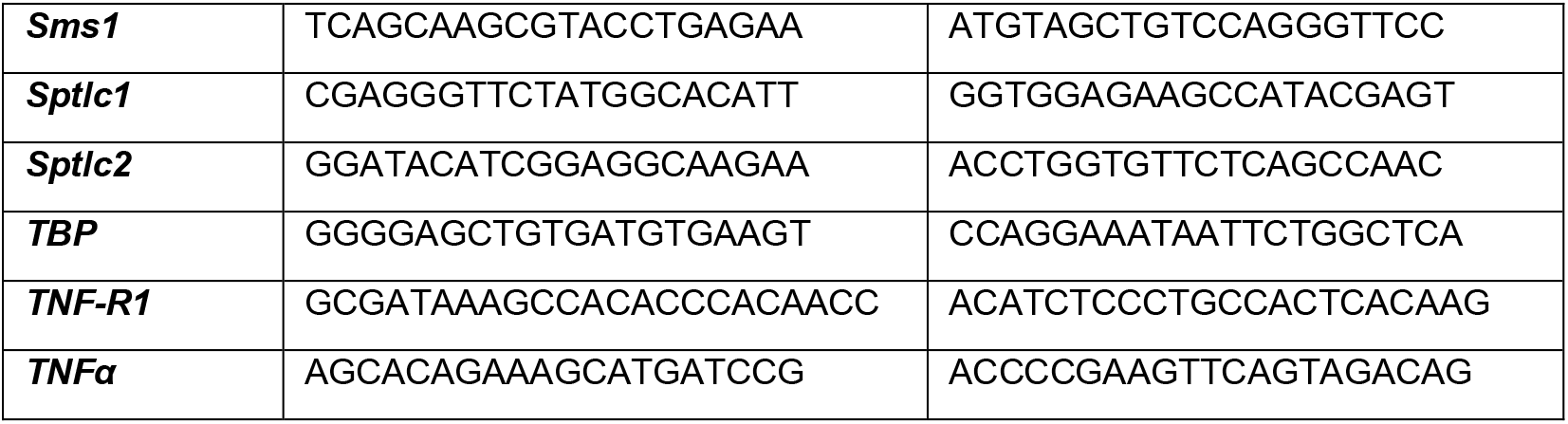
Gene expression analyzed by RT-qPCR and primer sequences.

## Acknowledgment

This project has received funding from the Innovative Medicines Initiative 2 Joint Undertaking under grant agreement No 115881 (RHAPSODY). This Joint Undertaking receives support from the European Union’s Horizon 2020 research and innovation program and EFPIA.

This study has also received funding from EUR G.E.N.E project (ANR-17-EURE-0013) and is part of the idEx project #ANR-18-IDEX-0001 of the Université de Paris, both funded by the French government under its Programme d’Investissements d’Avenir.

We acknowledge the technical platform Functional and Physiological Exploration platform (FPE) for body composition analysis (Université de Paris, BFA, UMR 8251 CNRS, Paris, France), the animal core facility “Buffon” of the University de Paris, Institut Jacques Monod, especially Laëtitia Pontoizeau, for animal husbandry and breeding.

We also acknowledge Dr. Fréderic Preitner and Anabela Rebelo Pimentel Da Costa from the Mouse Metabolic Facility in Center for Integrative genomics, University of Lausanne, Lausanne, Switzerland for measurement of energy content in faeces; Nicolas Sorhaindo from biochemical analyses facility in Inflammation research center, (Université de Paris, UMR 1149 Inserm, ERL CNRS 8252, Paris, France) for plasma biochemical analyses and Hermine Kakanakou and Sylvie Le Marchand from the genotyping and biochemical facility, Cordeliers research center, (Sorbonne Université, Paris, France) for mouse genotyping. We are grateful to the “Biogenouest Corsaire” core facility for mass spectrometry analyses.

We also acknowledge Amandine Gautier-Stein from Université Claude Bernard Lyon 1, (Université de Lyon, INSERM UMR-S1213, Lyon, France) for her scientific advice and protocols.

